# Generation of Self-Organising Macrovascular Constructs by Bioprinting human iPSC-Derived Mesodermal Progenitor Cells

**DOI:** 10.64898/2026.03.16.712040

**Authors:** Leyla E. Dogan, Nathaly A. Chicaiza-Cabezas, Florian Kleefeldt, Philipp Wörsdörfer, Jürgen Groll, Süleyman Ergün

## Abstract

Vascularization remains a major obstacle in tissue engineering. Here, we introduce a developmentally inspired bioprinting strategy to generate centimeter-scale, self-organising “mother vessel” constructs from iPSC-derived human mesodermal progenitor cells (hiMPCs). By systematically optimizing the bioink composition, we identified a formulation that combines high print fidelity, mechanical stability and cell compatibility within a single-step bioprinting process. Within the first week after printing, hiMPCs in the “mother vessel” constructs underwent spontaneous differentiation and morphogenesis, forming intima-, media-, and adventitia-like layers containing CD31⁺ endothelial, αSMA⁺ mural and CD34⁺/CD150⁺ progenitor cells. Remarkably, Iba1⁺ macrophage-like cells appeared despite their absence in the initial population, indicating intrinsic differentiation into both vascular and non-vascular lineages essential for angiogenesis, remodeling and tissue homeostasis. Surrounding the newly formed vessel wall-like structure was a broad, vascularized mesodermal tissue compartment that also contained the above-mentioned progenitors. Co-culture with prevascularized mesodermal organoids resulted in early structural interconnection of microvessels with the printed wall, representing a prerequisite for subsequent hierarchical vascular network formation. As a proof-of-concept, the mother vessel withstood controlled flow conditions in a bioreactor without detectable leakage, demonstrating its principal suitability for perfusion analyses. Together, these findings establish a biologically driven platform that bridges macro- and microvascularization. This may pave the way toward perfusable, vascularized larger tissue constructs, a major bottleneck in regenerative biofabrication.

## Introduction

A functional and well-organized vascular network is essential for maintaining tissue homeostasis, ensuring a continuous supply of oxygen and nutrients while removing carbon dioxide and other metabolic by-products. The vascular system forms a branching hierarchical network that ranges from large arteries and veins to smaller arterioles, capillaries and venules. Except for capillaries, all blood vessels share a characteristic triple-layered wall: the inner intima is formed by endothelial cells (ECs) and their basement membrane lining the vessel lumen, the media contains smooth muscle cells (SMCs) that mediate contractile functions and the outer adventitia harbors fibroblasts as well as stem and progenitor cells (Patan, 2004; Worsdorfer et al., 2017). Vascularization is essential for advancing 3D *in vitro* tissue models that are used to study human development, physiology and disease and may serve as tissue replacement in future. However, achieving proper vascularization remains one of the major challenges in bioengineering across tissues. Current approaches in vascular tissue engineering mostly combine adult vascular cell types such as ECs, SMCs and fibroblasts with supportive scaffold materials. These scaffolds range from decellularized native vessels (Conklin et al., 2002; Feiner et al., 2016) to synthetic and natural hydrogels (e.g. alginate, gelatin methacryloyl (GelMA), poly(ethylene glycol) diacrylate (PEGDA), collagen and fibrin) (Dogan et al., 2021; Gao et al., 2019; Gold et al., 2021; Peterson et al., 2014; Wang, 2019; Weinberg & Bell, 1986). The matrices are intended to provide a supportive environment for cell adhesion, survival and extracellular matrix formation that enables the fabrication of tubular constructs. However, most of the existing bioprinted vascular models published so far neither reproduce the multilayered vascular wall of native vessels nor a hierarchical network organization as observed *in vivo* that largely limits their translational relevance.

During embryonic development, *de novo* vessel formation, -termed vasculogenesis-, is driven by endothelial cells that differentiate from mesodermal progenitors and subsequently self-organize into an immature capillary plexus at intra- and extraembryonic sites (Risau, 1997). These processes are mainly governed by vascular endothelial growth factor A (VEGF-A). Likewise, the embryonic dorsal aorta, -initially formed as paired vessels that subsequently fuse into a single dorsal aorta-, develops through similar mechanisms and is initially composed solely of an endothelial tube. The entire vascular network then undergoes extensive remodeling, ultimately giving rise to a unified vascular system consisting of interconnected macro- and micro-vessels organized in a hierarchical manner.

Reproducing these morphogenetic processes *in vitro* by integrating bioprinting with the intrinsic self-organizing capacity of hiMPCs therefore represents a promising biologically inspired strategy to achieve structurally and functionally mature macrovessels in centimeter- and millimeter-scale that are hierarchically interconnected with engineered microvascular networks in tissues, e.g. organoids.

In line with this, we previously succeeded in generating mesodermal progenitors (hiMPCs) from human iPSCs that exhibited the capacity to differentiate *in vitro* into all cell types required for blood vessel formation, thereby eliminating the need for pre-differentiated vascular cells. Moreover, following extrusion-based bioprinting, these hiMPCs retained their multipotency, underwent spontaneous differentiation, and self-organized into hierarchically structured, blood vessel-like constructs that recapitulate key features of early embryonic vascular development (Dogan et al., 2021).

However, their intrinsic self-organizing capacity alone is not sufficient to form larger conduit vessels (>1mm inner diameter) that are required for functional perfusion or surgical integration with the host vasculature *in vivo*. To overcome this limitation, we employed 3D bioprinting to predefine millimeter-scale “mother vessels” designed to interconnect with the vascular networks of prevascularized surrounding tissues, such as organoids, and to serve as defined in- and outflow channels for these tissue constructs. To implement this biologically driven concept, we evaluated different hydrogel formulations for their suitability in extrusion bioprinting and vascular morphogenesis. We were able to identify a blend matrix that provided a balance between printability, mechanical stability and cellular compatibility. Using this optimized formulation, we successfully printed centimeter-scale “mother vessel” constructs that exhibited early vascular layering and were suitable for initial perfusion experiments. Furthermore, direct co-culture with prevascularized organoids demonstrated early structural interconnection with the “mother vessel” representing a first step toward assembling larger tissues through a modular building-block approach. This single-step, developmentally guided strategy offers a simple approach to generate macro-scale vascular tissues that recapitulate key aspects of embryonic vasculogenesis that will facilitate to bridge the gap between large host vessels and microvascular networks within engineered tissues. Moreover, a thick matrix layer containing hiMPCs at various stages of differentiation, which display immunophenotypic markers characteristic of mesodermal, endothelial, and hematopoietic lineages surrounds the “mother vessel”.

## Material and Methods

### Cell culture and differentiation

Human induced pluripotent stem cells (hiPSCs) were generated from commercially available human dermal fibroblasts using the hSTEMCCA lentiviral reprogramming vector as described previously (Kwok et al., 2018; Sommer et al., 2009). hiPSCs were maintained on Matrigel^®^-coated culture dishes (hESC-qualified matrix, 354277, Corning, New York, USA) in StemMACS iPS Brew medium (130-107-086, Miltenyi Biotec, Bergisch Gladbach, Germany) with daily medium exchange. For routine passaging, cultures at approximately 80% confluence were dissociated into single cells using StemPro Accutase (A6964, Merck, Temecula, USA) for 5 min at 37 °C. The cells were reseeded onto Matrigel^®^-coated dishes in StemMACS medium supplemented with 10 nM thiazovivin (TZ, T9753, LC Labs, Woburn, MA, USA) to enhance cell survival and attachment.

HiPSCs were differentiated into mesodermal progenitor cells (hiMPCs) using a previously established differentiation protocol (Wörsdörfer et al., 2019). Confluent hiPSC cultures were dissociated with Accutase, counted and seeded on Matrigel^®^-coated six-well plates (3.5×10^5^ cells/cm^2^). Cells were maintained for 24 h in StemMACS iPS-Brew medium supplemented with 10 nM TZ. The medium was then replaced with mesodermal induction medium (Advanced DMEM/F-12, 126-34010, Thermo Fisher Scientific) containing 0.2 mM L-glutamine (G7313, Merck), 60 µg/mL ascorbic acid (A4544, Sigma-Aldrich, St. Louis, USA), 10 µM CHIR99021 (SML1046, Merck), 25 ng/mL BMP4 (PHC9534, Thermo Fisher Scientific) for three days at 37 °C and 5% CO_2_ with daily medium replacement. After induction, cells were dissociated again with Accutase and collected for bioprinting. Printed “mother vessel” constructs were transferred to culture flasks and maintained in vascular growth medium (Advanced DMEM/F-12 supplemented with 5% heat-inactivated FCS (S00H81000C, Biowest, Karlruhe, Germany), 1% penicillin/streptomycin (P0781, Merck), 10 nM TZ, 0.2 mM L-glutamine, 60 µg/mL ascorbic acid and 60 ng/mL VEGF-A (10020, Thermo Fisher Scientific)) at 37 °C and 5% CO_2_.

3D prevascularized mesodermal organoids were generated as described in a previously published protocol (Wörsdörfer et al., 2019). However, instead of using 96-well plates, custom made agarose molds were utilized. These honeycomb-shaped microwell molds were generated using a 2% (w/v) agarose solution (840004, Biozym, Hessisch Oldendorf, Germany) prepared in filtered Ampuwa^®^ (1080181, Freseneus Kabi, Bad Homburg, Germany). The hot agarose solution was poured over 3D-printed negative molds placed in wells of a 12-well plate and allowed to solidify for 1h at RT. Subsequently, the agarose was gently separated from the 3D-printed negatives, resulting in molds with 159 microwells, each approximately 700 µm in diameter. The agarose molds were filled with DMEM supplemented with 1% penicillin/streptomycin and sterilized under UV light O/N. Prior to seeding, DMEM was removed and replaced by MACS TZ medium supplemented with TZ (1:4000). 4×10^4^ cells of a hiPSCs single-cell suspension were seeded into each mold. Cells were cultured for 1 day in MACS TZ medium, followed by 3 days in mesoderm induction medium (MIM) as described above. On day 4, the medium was replaced with vascular growth medium, in which the organoids were maintained until they reached the printable size (approximately 300µm) at culture day 7.

## Bioink preparation

### Material synthesis

Porcine GelMA and piscine GelMA were synthesized following a protocol published previously (Loessner et al., 2016) Briefly, gelatin from porcine skin or cold-water fish skin (G7041 and G1890, Sigma-Aldrich) was dissolved in phosphate-buffered saline (1x PBS, D1408, Sigma-Aldrich) at a concentration of 10% w/v at 37 °C. Methacrylic anhydride (276685, Sigma-Aldrich) was added dropwise at 60% (v/v) under continuous stirring and the reaction was allowed to proceed for 1 h at 37 °C. Unreacted methacrylic anhydride was removed via centrifugation and the supernatant was diluted twofold with prewarmed Milli-Q water (37 °C). The solution was transferred to a dialysis membrane (MWCO 3.5 kDa) and dialyzed against Milli-Q water for three days. The resulting GelMA solution was then lyophilized and stored at −20°C until further use.

### GelMA preparation

Porcine GelMA (5% w/w) was dissolved in PBS under magnetic stirring (200 rpm) at 47 °C. After complete dissolution, lithium phenyl-2,4,6-trimethylbenzoylphosphinate (LAP, 0.1% w/v; 900889, Sigma-Aldrich) was added and the solution was stirred for an additional 10 min. The resulting GelMA-LAP solution was maintained in a 37 °C water bath until further use. For bioink preparation, hiMPCs were detached, counted and resuspended in the prewarmed GelMA solution at a final concentration of 2×10^7^cells/mL.

### GelMA+Col I preparation

Collagen type I (Col I, 4.7 mg/mL; rat tail, 08-115, Merck) was neutralized according to the manufacturer’s instructions. The neutralized Col I was blended with 5% (w/w) porcine GelMA at a 1:4 ratio using a positive displacement pipette, resulting in a final composition of 3.75% (w/w) GelMA and 0.68 mg/mL Col I. For bioink preparation, hiMPCs were detached, counted and resuspended in the neutralized collagen solution. The cell-collagen suspension was then combined with 5% porcine GelMA in the same 1:4 ratio to yield the final bioink containing 2×10^7^ cells/mL.

### FGX preparation

Piscine GelMA (3% w/w) and porcine GelMA (2% w/w) were dissolved in vascular growth medium under magnetic stirring (200 rpm) at 47 °C. After complete dissolution, LAP (0.1% w/v) and xanthan gum (0.5% w/v) (06-003, Cosphaderm X34, Cosphatec, Hamburg, Germany) were added. The solution was stirred for 45 min (100 rpm, 47 °C) to ensure homogeneity. The resulting blend (FGX) was maintained in a 37 °C water bath until cell addition. For bioink preparation, hiMPCs were detached, counted and resuspended in the prewarmed FGX blend at a final concentration of 2×10^7^ cells/mL.

### FGXC preparation

Piscine GelMA (6% w/w) and porcine GelMA (4% w/w) were dissolved in vascular growth medium under magnetic stirring (200 rpm) at 47 °C. After complete dissolution, LAP (0.2% w/v) and xanthan gum (1% w/v) were added and the mixture was stirred for 45 min at 100 rpm and 47 °C to ensure homogeneity. The resulting blend (2× FGX) was subsequently mixed at a 1:1 ratio with 1% Fibercoll-Flex N^®^ bioink (500069016, Viscofan Bioengineering, Weinheim, Germany), prepared according to the manufacturer’s instructions. The final composite blend (FGXC) contained 3% piscine GelMA, 2% porcine GelMA, 0.5% xanthan gum, 0.1% LAP and 0.5% fibrillar Col I. For bioink preparation, hiMPCs were detached, counted and resuspended in the prewarmed FGXC blend at a final concentration of 2×10^7^ cells/mL.

### Support bath preparation

Hyaluronic acid sodium salt (0.2% w/v, 80-100 kDa, FH63427, Biosynth, Staad, Switzerland) was dissolved in HEPES buffer (H4034, Sigma-Aldrich, pH 7.4, 10 mM) under magnetic stirring (500 rpm, RT). After complete dissolution, the solution was passed through a 0.2 µm filter. UV-sterilized xanthan gum (1% w/v) was added, and the mixture was stirred (700 rpm, RT) until a homogeneous solution was obtained.

### Rheological characterization

Rheological measurements were performed using an MCR 702 MultiDrive rotational rheometer (Anton Paar, Graz, Austria) equipped with a 25 mm parallel-plate geometry at 25 °C. For bioink characterization, shear rate sweep tests were carried out to determine the apparent viscosity by increasing the shear rate from 0.001 to 100/s over 300s. Temperature-dependent viscoelastic behavior was analyzed by temperature sweep measurements of complex viscosity (η*), storage modulus (G′) and loss modulus (G″). The temperature varied at a rate of 0.01°C/s in four steps: 1) heating from 15 to 45°C, 2) isothermal hold at 45°C, 3) cooling from 45 to 15°C and 4) isothermal hold at 15°C. For the embedding medium, amplitude sweep measurements were conducted over a strain range of 0.01-1000% at a constant angular frequency of 10rad/s. Thixotropic behavior and stress recovery were evaluated in a five-step test sequence: 1) low shear stress for 60s, 2) high shear stress for 10s, 3) low shear stress for 60s, 4) high shear stress for 10s and 5) low shear stress for 60s.

### Embedded 3D Bioprinting

Tubular constructs were designed in Autodesk Fusion 360 (Autodesk, San Francisco, CA, USA) as cylinders with a diameter of 4.6 mm and heights of 5, 10 or 20 mm. The resulting CAD models were exported and sliced in Slic3r using a 0% infill and a layer height of 0.2 mm. Cell-laden bioinks were transferred into 3 ml syringe barrels (Nordson), sealed with tip caps and centrifuged at 1200 rpm for 2 min at RT to remove air bubbles. Subsequently, the tip caps were replaced with 22G (1 inch, 0.41 mm inner diameter) or 20G (1 inch, 0.61 mm inner diameter) needles and the prepared syringes were mounted into the printhead of a BIO X bioprinter (CELLINK, Gothenburg, Sweden). The specific printing parameters used for each bioink formulation are summarized in Table 1.

**Table 1:** Printing Parameters for different bioink formulations.

| Bioink | Pressure (kPa) | Speed (mm/min) | Needle gauge | Temperature (°C) |
| --- | --- | --- | --- | --- |
| GelMA | 15-25 | 300 | 22 G | 18-24 |
| GelMA+Col I | 15-20 | 300 | 22 G | 18-24 |
| FGX | 4-5 | 300 | 22 G | RT |
| FGXC | 6-7 | 360 | 20 G | RT |

For 5 and 10 mm tubular constructs, 12-well plates were filled with the support bath, and one tube was printed per well. The entire plate was then exposed to 405 nm light (54 mW cm⁻²) for 3 min to induce photopolymerization. After crosslinking, the printed tubes were carefully removed from the support bath, rinsed in PBS to remove residual xanthan gum and transferred into cell culture medium. For 20 mm constructs, glass vials were filled with the support bath, and the longer tubes were printed directly inside the vials. Crosslinking and washing were performed as described above.

### Printability characterization

The inks were prepared and kept at RT for at least 20 min before printing. The printability tests were performed using a 22G needle nozzle, 0.25 inch. A filament fusion test (FFT) was used to define the category of the ink, according to Lamberger et al.(Lamberger et al., 2024) and a semi-quantitative evaluation based on the pore geometry (Pr) was used to characterize printed grids, according to Ouyang et al. (Ouyang et al., 2016).

### Mechanical characterization

Compression testing was performed using a dynamic mechanical tester (ElectroForce 5500, Bose; TA Instruments, New Castle, DE, USA). Hydrogel samples were extruded into molds, crosslinked under 405 nm light (3 min, 54 mW cm^-2^) and tested using a 250g load cell. The Young’s modulus was determined from the slope of the linear region of the stress-strain curve within the viscoelastic range.

### Cryo-scanning electron microscopy

The microstructure of the hydrogels was examined using cryo-scanning electron microscopy (cryo-SEM). Samples were extruded, crosslinked and analyzed immediately after preparation. In addition, FGXC samples were incubated for 2 and 6 days prior to analysis to assess structural alterations over time. For cryo-SEM preparation, hydrogel specimens were placed between aluminum plates (3 mm diameter) and rapidly frozen in slushed liquid nitrogen at −210 °C. The frozen samples were transferred using an EM VCT100 cryo-shuttle (Leica Microsystems) into an ACE 400 sputter coater (Leica Microsystems) at −140 °C. After removing one aluminum plate, the exposed fracture surface was etched for 15 min at −85 °C under vacuum (<1×10⁻³ mbar). The surface was then sputter-coated with 3 nm platinum and transferred to a Crossbeam 340 SEM (Zeiss). Imaging was performed at −155 °C with an accelerating voltage of 8 kV.

### Cell viability

To assess cell viability during culture, live/dead staining was performed using Calcein AM (2 µM; C3099, Life Technologies) and Ethidium Homodimer-1 (EthD-1, 4 µM; E1169, Life Technologies) according to the manufacturer’s instructions. Printed tubular constructs from independent experiments were analyzed at days 1, 3 and 7. At each time point, three constructs were rinsed three times 5 min with PBS on a rocker to remove residual medium. Samples were then fully immersed in freshly prepared staining solution and incubated at 37°C for 1 h. Following incubation, the staining solution was replaced with PBS and the constructs were washed three times for 15 min each on a rocker. Fluorescence imaging was acquired using a confocal laser scanning microscope (Nikon Eclipse Ti with Nikon NIS-Elements software version 4.13.05, Nikon, Tokyo, Japan). Images were analyzed using FiJi software to quantify live and dead cell fractions. Data visualization and statistical analysis (Student T test) were performed using Prism5 Software (GraphPad, LaJolla, CA, USA).

### Immunofluorescence

To evaluate the cellular phenotype prior to bioprinting, cells maintained in 2D culture were fixed with 4% paraformaldehyde (PFA; P6148, AppliChem, Darmstadt, Germany) for 15 min at RT. After fixation, samples were blocked for 1 h at RT in 4% bovine serum albumin (BSA; A1391, AppliChem) blocking buffer solution. For intracellular markers, 0.1% Triton X-100 (T8787, Sigma–Aldrich) was added to the blocking buffer to permeabilize cell membranes, whereas surface markers were stained without detergent. Cells were then incubated O/N at 4 °C with primary antibodies diluted in blocking buffer. The following antibodies were used: OCT4 (sc-5279, Santa Cruz Biotech, Dallas, TX, USA), SOX2 (MAB2018, Santa Cruz Biotech), Brachyury/T (AF2085, Santa Cruz Biotech) and VEGFR-2 (sc-48161, Santa Cruz Biotech). After washing three times with PBS, cells were incubated for 1h at RT with Cy2- or Cy3-conjugated secondary antibodies (Dianova, Hamburg, Germany) diluted in PBS. Fluorescence imaging was performed using a Biorevo fluorescence microscope (Keyence, Osaka, Japan). The analyses confirmed that hiMPCs expressed mesodermal progenitor markers Brachyury and VEGFR-2 **(Fig.1 B2-B3).**

**Figure 1:**
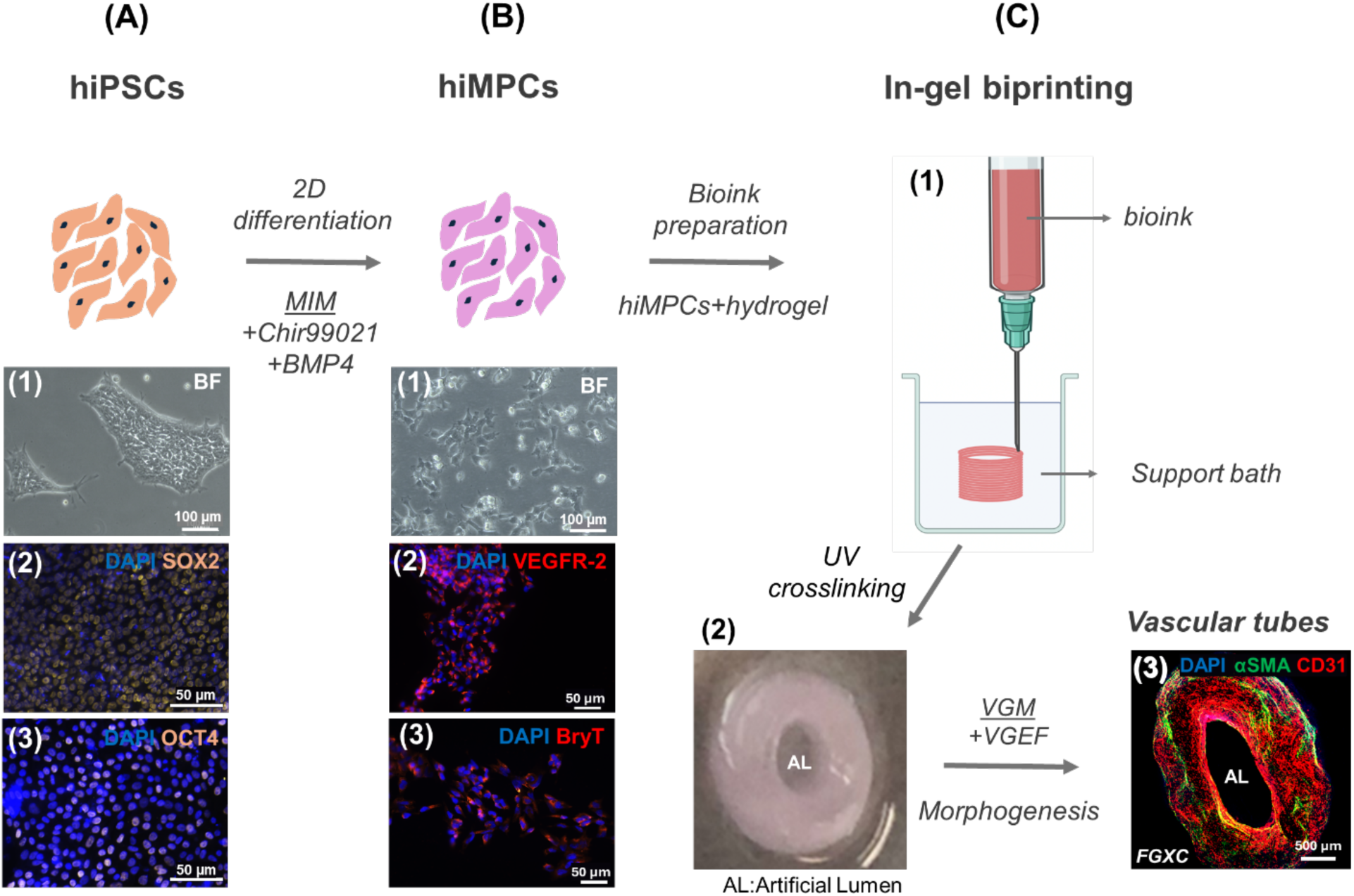
Workflow for generating hiMPCs form hiPSCs and printing of hiMPCs within hydrogel to produce vascularized tubular structures. **(A1)** and **(B1)** Phase contrast images of hiPSCs and HiMPCs. **(A1)** hiPSCs and **(B1)** hiMPCs cultured in standard 2D conditions. **(A2 & A3)** Immunofluorescence images showing hiPCS stained for SOX2 and OCT4 (both in yellow). (B2 & B3) hiMPCs stained for VEGFR-2 and BryT (both in red). **(C1)** Schematic representation of the in-gel printing-embedded bioprinting-approach. **(C2)** Printed and crosslinked FGXC tube. **(C3)** Maximum intensity projection of z-stack images from a cleared printed tube at day 12. The endothelization around the artificial lumen and endothelial network within the tube wall is shown by CD31^+^ staining (red). αSMA^+^ cells are shown in green.

For histological analysis, bioprinted tubular constructs were washed three times in PBS (10 min each) and fixed O/N in freshly prepared 4% PFA at 4 °C. Fixed samples were washed in PBS (3 × 30 min) to remove residual fixative and stored in 70% ethanol (S642, Nordbrand, Darmstadt, Germany) until paraffin (1.116092504, CarlRoth, Darmstadt, Germany) embedding. Sections (10 µm) were deparaffinized, rehydrated and stained with hematoxylin (50837, Chroma, Darmstadt, Germany) and eosin (A0822, Chroma, Darmstadt, Germany) (H&E) or processed to visualize collagenous matrix components. For immunofluorescence on paraffin sections, antigen retrieval was performed in 10 mM sodium citrate buffer (pH 6.0). Sections were blocked for 1h at RT in 4% BSA. For intracellular markers, 0.1% Triton X-100 was added. After blocking, sections were incubated overnight at 4 °C with primary antibodies diluted in blocking buffer. The following primary antibodies were used: CD31 (M0823, DAKO, Agilent, Santa Clara, CA, USA), αSMA (ab5694, Abcam, Cambridgeshire, UK), CD34 (130-105-830, Miltenyi Biotech), CD150 (PA5-21123, Invitrogen), CD44 (103002, BioLegend, San Diego, CA, USA), Collagen IV (ab6586, Abcam), CD45 (13-9457-82, Invitrogen) and IBA1 (019-19741, Wako/Fuji Film, Neuss, Germany).

After washing three times in PBS, sections were incubated for 1h at RT with fluorophore-conjugated secondary antibodies (Dianova, Hamburg, Germany) goat anti-rabbit Cy2 (111-225-144), goat anti-rabbit Cy3 (111-165-003), goat anti-mouse Cy2 (115-225-146), goat anti-mouse Cy3 (115-165-003) and goat anti-rat Cy5 (112-175-143). Nuclei were counterstained with DAPI (2360276, Merck). Stained sections were imaged using a Nikon Eclipse Ti confocal laser scanning microscope (Nikon).

For vibratome sections, printed tubular constructs were washed three times with PBS (10 min each) on a shaker and subsequently fixed in 4% PFA (in 0.1M PBS O/N at 4 °C). Excess fixative was removed by washing the samples three times with PBS for 30 min each. The fixed constructs were embedded in warm 1% agarose prepared in distilled water (dH_2_O). After solidification, the agarose blocks were mounted onto the cutting platform using superglue and sectioned (300µm) with a vibratome (VT1000S, Leica Microsystems, Germany). Blocking was performed for 2h at RT, followed by overnight incubation with primary antibodies at 4 °C and 3h incubation with Cy2- or Cy3-conjugated secondary antibodies at RT in PBS.

### H&E staining

Following deparaffinization slides were immersed in hematoxylin solution (Chroma^®^, 50837, Waldeck GmbH, Münster, Germany) for 10 min, rinsed briefly with distilled water and then washed for 10 min under running tap water. After a final rinse with distilled water, cytoplasmic counterstaining was performed with eosin for 10 min. Subsequently, slides were dehydrated through graded ethanol (96% for 2 min, followed by two changes of 99% for 5 min each) and cleared twice in 99.9% xylene (University of Würzburg, Würzburg, Germany) for 5 min. Finally, specimens were mounted with DePeX mounting medium (10236276001, SERVA Electrophoresis GmbH, Heidelberg, Germany).

### Tissue Clearing

To visualize vascular network formation within the printed constructs, whole-mount immunofluorescence staining was combined with ethyl cinnamate-based tissue clearing as described previously (Worsdorfer et al., 2020). Endothelial cells (ECs) and peri-endothelial cells (PECs) were labeled using primary antibodies against CD31 and αSMA, respectively. Fluorescence was visualized with Cy2- and Cy3- conjugated secondary antibodies. Confocal imaging was performed as described above.

### Perfusion of the mother vessel

We designed and fabricated a prototype perfusion chamber to mimic physiological flow conditions by enabling controlled perfusion through 3D-bioprinted tubular structures. The chamber allows parallel perfusion of two tubular constructs within a shared chamber geometry, enabling simultaneous flow through two vessels in a defined and aligned orientation. The chamber accommodates the printed constructs within a rectangular compartment (22 mm × 22 mm × 8 mm; L × W × H) and is perfused via Luer-lock ports after assembly. Sealing between the upper and lower half-shells is achieved using a two-component silicone (SF33, silicone fabric). The printed perfusion chamber is reproducible, readily customizable to experimental requirements, and biocompatible.

The upper and lower half-shells that jointly form the perfusion chamber were designed using 3D CAD software (SolidWorks). The corresponding STL files were transferred to a 3D printer (Elegoo Saturn 4 Ultra, 16K) for fabrication. Both components were printed using a biocompatible resin based on BioMed Clear (LiQcreate, The Netherlands) and subsequently post-cured under 405 nm UV light for 20 min. After printing, the components were removed from the build platform and washed in 96% ethanol (MEK, Nordbrand) to remove residual resin.

To generate a closed medium reservoir and enhance optical clarity, a 1-mm glass plate was bonded to the top surface of the upper half-shell and to the bottom surface of the lower half-shell. The two half-shells were then aligned and secured at the corners using small screws. Lateral interfaces were sealed with plastic Luer-lock adapters (Borem; Luer-lock to 1/8″ tubing). Phosphate-buffered saline (PBS) was injected through one adapter using a syringe, and the fully assembled chamber was filled and incubated overnight to assess leakage. No leakage was observed.

Prior to assembly, all components of the perfusion setup were sterilized 3x 30min by immersion in 70% ethanol. The parts were subsequently removed from ethanol, rinsed twice in sterile PBS for 10 min each, air-dried under a sterile bench for 1 h, and post-cured under UV light for 15 min on each side. Following UV curing, day-7 bioprinted tubular constructs were assembled into the perfusion chamber and perfused under slow pulsatile flow (1 mL/min) using oxygenated basal culture medium delivered by a peristaltic pump system (Golander B100, adjustable flow mode and rate; Golander GmbH, Bonn, Germany).

## Results and Discussion

After confirming the pluripotency of the hiPSC line via immunofluorescence (IF) staining for markers such as SOX2 and OCT4 **(Fig. 1A2-A3)**, the cells were differentiated into hiMPCs using a previously established 2D-culture protocol (Wörsdörfer et al., 2019). IF staining for mesodermal progenitor markers, including VEGFR-2 and Brachyury, confirmed successful differentiation **(Fig. 1B2–B3).** hiMPCs were selected as cell sources for bioink formulation, as they have the ability to differentiate into all cell types of the vascular wall and form hierarchically organized, perfusable vessel-like structures (Dogan et al., 2021). For each experimental batch, hiMPCs were freshly induced shortly before printing. The bioink was prepared by evenly suspending the cells at a density of 2×10^7^ cells per milliliter of hydrogel. Throughout this study, we employed embedding bioprinting **(Fig. 1C1)** to fabricate large-scale tubular structures by depositing bioinks into a XG-based support bath. Following printing, the structures were stabilized via UV crosslinking and subsequently cultured under static conditions for a defined period **(Fig.1C).**

### Bioink characterization and optimization

To develop a bioink that combines both reliable printability and the capacity to support vascular morphogenesis, several candidate hydrogels were initially evaluated. Preliminary experiments with alginate- and fibrin-based matrices revealed substantial limitations, as neither material allowed the cells to undergo morphogenetic processes required for vessel wall formation.

We therefore focused on gelatin methacryloyl (GelMA) as a versatile base matrix with tunable physical and biological properties. However, porcine GelMA alone was unable to provide the thermal stability, rheological control and mechanical robustness required for the fabrication of large “mother vessel” constructs. To overcome these limitations, we systematically optimized the formulation by incorporating defined additives with complementary functions: Col I to enhance cell adhesion and mimic native extracellular matrix composition, piscine GelMA to improve thermal handling during the printing process due to its lower gelling temperature, xanthan gum to tune viscosity and enable shear-thinning behavior and fibrillar Col I to reinforce the mechanical stability and homogeneity of the matrix.

Four GelMA-based blends of increasing compositional complexity were compared: 1) GelMA without any additives as baseline, 2) GelMA + Col I, 3) FGX (piscine + porcine GelMA + xanthan gum) and 4) FGXC (FGX + fibrillar Col I). Each formulation was systematically characterized for rheological behavior, printability, mechanical performance and microstructural organization **(Fig. 2).** Rheological measurements revealed distinct viscoelastic profiles of the tested formulations. Temperature-sweep tests (15-45°C) showed that both FGX and FGXC maintained stable storage (G′) and loss (G″) moduli across the full range, indicating temperature-stable viscoelastic behavior and suitability for bioprinting under room temperature (RT) conditions. In contrast, GelMA and GelMA + Col I exhibited sharp thermal transitions around 30 °C, leading to unstable flow behavior and reduced process reliability. Shear-rate sweeps (0.001-100/s) further demonstrated pronounced shear-thinning for FGX and FGXC, enabling consistent extrusion at low pressures (< 15 kPa).

**Figure 2:**
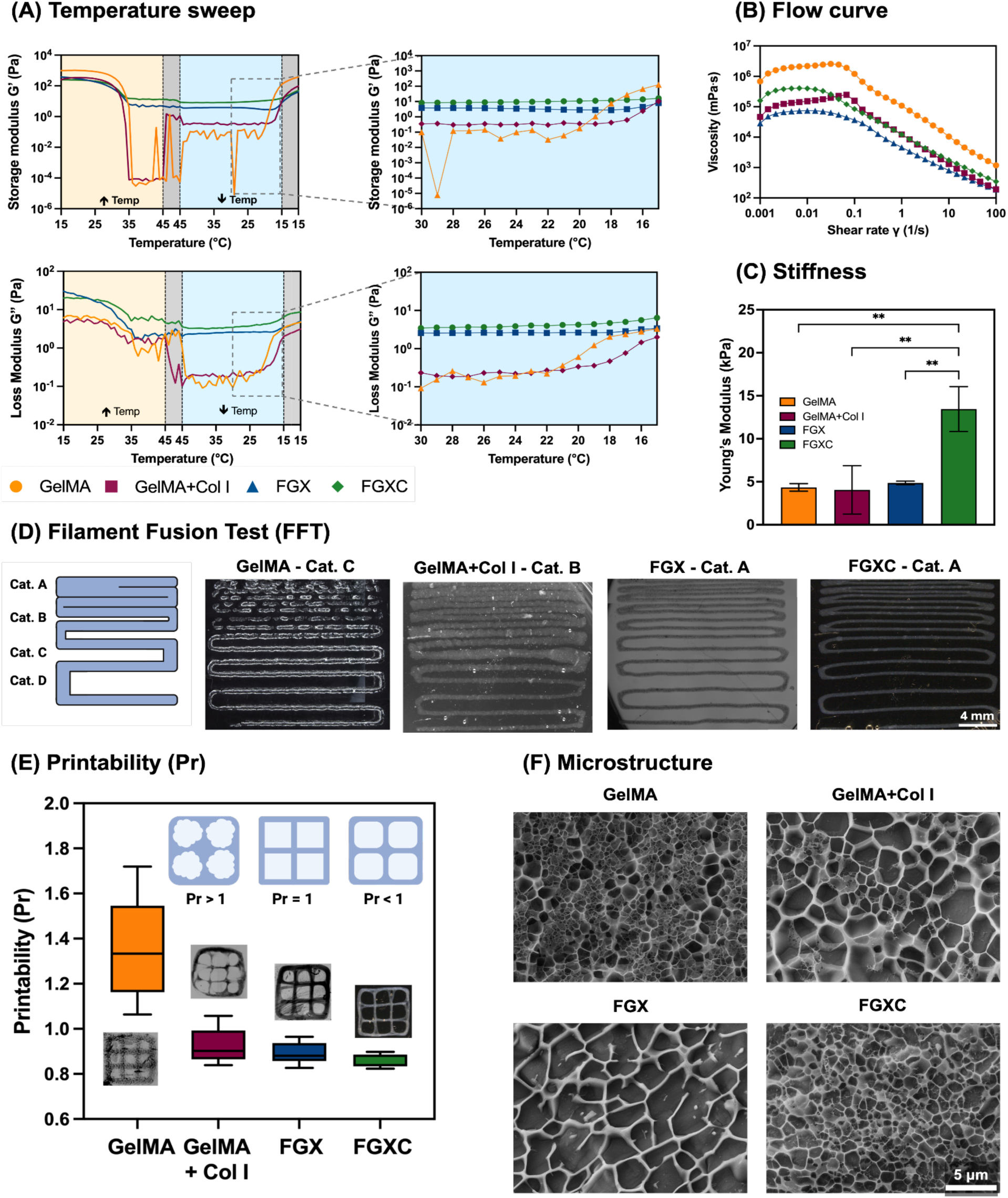
Rheological, printability, mechanical, and microstructural characterization of the tested hydrogel formulations. **(A)** Temperature sweep analysis showing storage modulus (G′) and loss modulus (G″) as a function of temperature prior to crosslinking. **(B)** Flow curves depicting viscosity as a function of shear rate for the non-crosslinked bioinks. **(C)** Compressive Young’s modulus of crosslinked hydrogels. **(D)** Filament fusion test (FFT) used to assess print resolution and categorize bioinks according to Lamberger et al.(Lamberger et al., 2024) **(E)** Quantitative printability index (Pr) analysis based on pore geometry according to Ouyang et al.(Ouyang et al., 2016). **(F)** Microstructural analysis of the crosslinked hydrogels by cryo-SEM.

Printability was initially determined using the filament-fusion test (FFT) and a printability index (Pr). GelMA (Pr = 1.35 ± 0.22, Cat C) and GelMA + Col I (Pr = 0.92 ± 0.07, Cat B) exhibited irregular pore geometry and intermittent filament flow, occasionally due to collagen aggregation. In contrast, FGX (Pr = 0.89 ± 0.05, Cat A) and FGXC (Pr = 0.87 ± 0.03, Cat A) yielded well-defined, continuous filaments with uniform deposition and minimal over- or under-extrusion.

Mechanical compression testing showed that the combined addition of xanthan gum and fibrillar Col I markedly increased the mechanical stability of the hydrogels. The corresponding Young’s moduli were 4.3 ± 0.4 kPa (GelMA), 5.2 ± 0.5 kPa (GelMA + Col I), 4.9 ± 0.2 kPa (FGX) and 13.4 ± 2.6 kPa (FGXC). Thereby, FGXC was in the mechanical range of soft connective tissue (Handorf et al., 2015). Cryo-SEM analysis further confirmed distinct microstructural differences among the tested hydrogels. FGXC exhibited a dense, homogeneous fibrillar network with uniformly distributed pores, whereas GelMA, GelMA + Col I and FGX were more heterogeneous with variable pore size and network density, consistent with their lower printability index and stiffness **(Fig. 2F).** Hence, the incorporation of cold-water-fish-derived GelMA improved thermal stability and printability, while fibrillar Col I enhanced mechanical strength and homogeneity of matrix organization. Together, these findings identify FGXC as the optimal bioink formulation, as it combines temperature stability, shear-thinning rheology and mechanical robustness with high print fidelity.

### Embedded 3D Bioprinting in an XG-HA Support Bath

For the direct bioprinting of free-standing tubular constructs, cell-laden bioinks were extruded into a viscoelastic support bath composed of xanthan and hyaluronic acid in HEPES buffer (see Methods). The support bath provided transient mechanical stabilization during extrusion. This enabled the use of soft, low-viscosity hydrogels that would otherwise collapse under their own weight. Moreover, the viscoelastic behavior of the XG-HA matrix created gentle shear conditions that preserved cell integrity and supported morphogenetic cell behavior during and after printing. The support bath exhibited self-healing, fluid-like behavior under applied stress and pronounced shear-thinning, as confirmed by stress-recovery, flow-curve and frequency-sweep analyses **(Suppl. Fig. S1C)**. Thereby, the support bath not only ensured temporary mechanical support during the printing process but also enabled the gentle release of the vascular constructs after crosslinking, maintaining structural integrity (Becker et al., 2023; Brunel et al., 2022; Lai & Meagher, 2024; Patricio et al., 2020; Tan et al., 2021).

Using the optimized FGXC formulation, tubular structures were printed at RT under low extrusion pressure (< 20 kPa) through 20–22 G needles (20 G for FGXC). Constructs were photopolymerized (405 nm, 3 min; 54 mW/cm²) and then gently released from the support bath by rinsing. This removed residual XG without affecting the printed geometry and resulted in self-standing tubes with interlayer fusion, homogeneous wall structure and tunable dimensions (up to 1.5–2 cm length, 3.5–4.5 mm inner lumen diameter) **(Fig. 4A1 & A2).**

In parallel, the biological evaluation of hiMPC-laden constructs revealed distinct morphogenetic patterns across the different bioink formulations **(Fig. 3A–D).** All formulations maintained high cell viability, with stable tube geometry and the formation of a continuous cell–cell layer around the artificial lumen. To improve cell adhesion and mimic the extracellular matrix composition of native vascular tissue, Col I was incorporated into GelMA. Both GelMA and GelMA + Col I supported early endothelial alignment. However, most cells accumulated along the tube borders, reflecting heterogeneous matrix density and uneven cell distribution within the hydrogel. To further enhance temperature stability and extrusion consistency, cold-water-fish-derived GelMA was introduced, and xanthan gum was added to tune viscosity and promote shear-thinning behavior during printing. The FGX formulation improved cell retention and uniform distribution throughout the tube wall and promoted cord-like endothelial structures. Finally, supplementing the FGX blend with fibrillar Col I provided additional mechanical reinforcement and improved matrix homogeneity. The optimized FGXC bioink combined these features, enabling homogeneous cell localization, rapid elongation and the formation of interconnected CD31^+^/αSMA^+^ vascular-like networks within seven days of culture **(Fig.3D).** Over time, the gradual release of XG increased matrix microporosity (**Suppl. Fig. S2)** and facilitated directed cell migration and network formation, underscoring the biocompatibility and morphogenetic potential of the FGXC system **(Fig.3D).**

**Figure 3:**
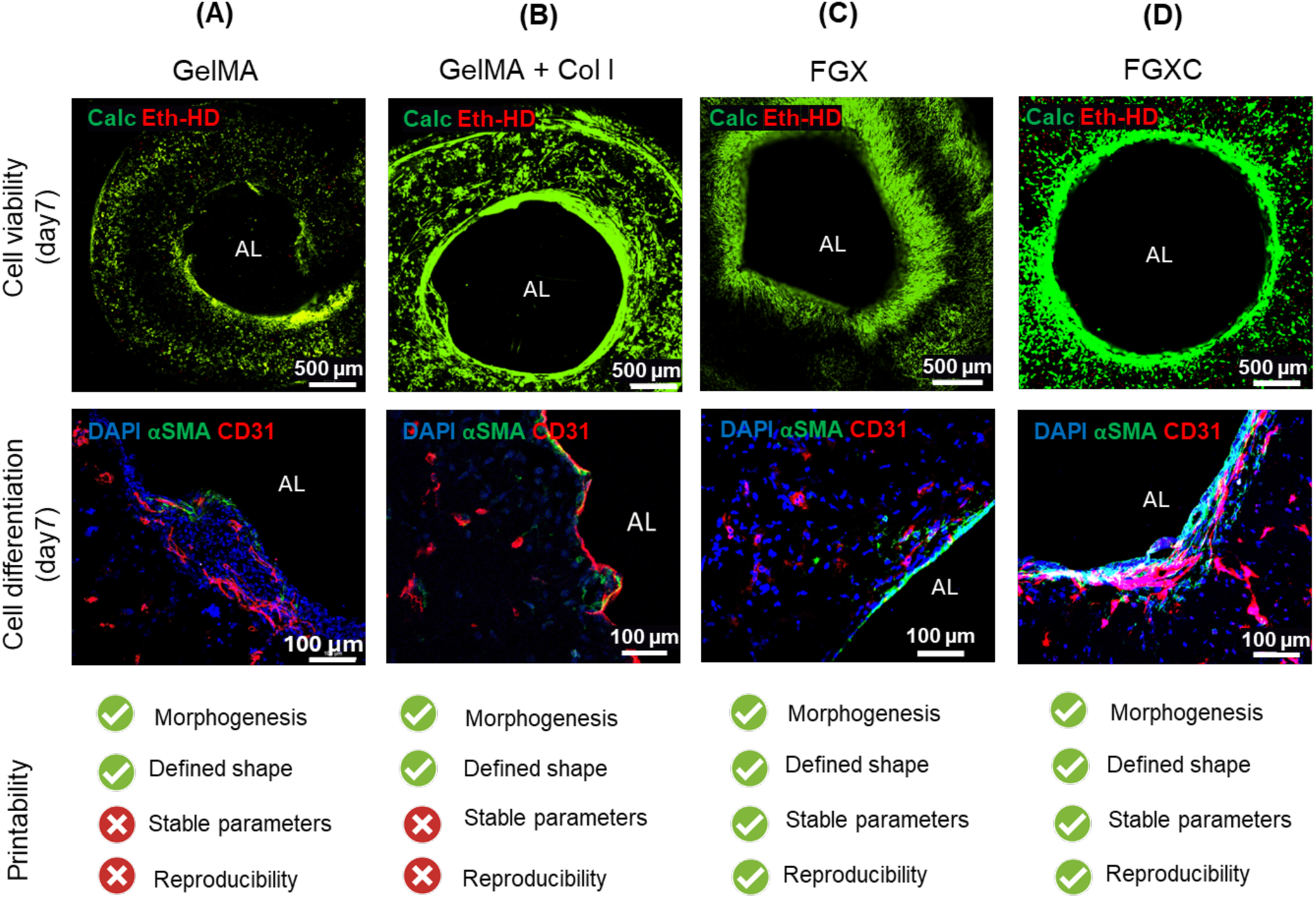
Morphogenic processes and printability of hiMPCs-laden bioinks. **Upper panel:** Representative fluorescence viability images at d7 (Calcein AM/EthD-1; initial cell density 2×10^7^ cells/ml) for the different hydrogel formulations. Under all conditions, the printed tubes remained structurally stable, exhibited high cell viability and displayed a clearly defined, continuous cellular layer that surrounded the artificial lumen (AL). **Middle panel:** Vascular morphogenesis within the printed constructs. hiMPCs differentiated into CD31^+^ endothelial cells (ECs; red) and αSMA^+^ pericyte-like cells (PECs; green), followed by spatial organization and the formation of vascular-like structures. While all hydrogel formulations supported cell adhesion and lineage specification, the degree of organized vascular differentiation varied between formulations. Cell nuclei are counterstained using DAPI. **Printability assessment:** Printability was evaluated based on shape fidelity of the printed tubes, stability of printing parameters and reproducibility of the resulting tubular structures.

### Vascular Morphogenesis and Vessel Wall Formation

Following confirmation of high post-printing viability **(Fig. 4C)**, vascular morphogenesis and vessel wall organization were examined within the bioprinted FGXC “mother vessels”. To assess temporal changes in vascular development, constructs were cultured under static conditions for up to 14 days and analyzed histologically.

**Figure 4:**
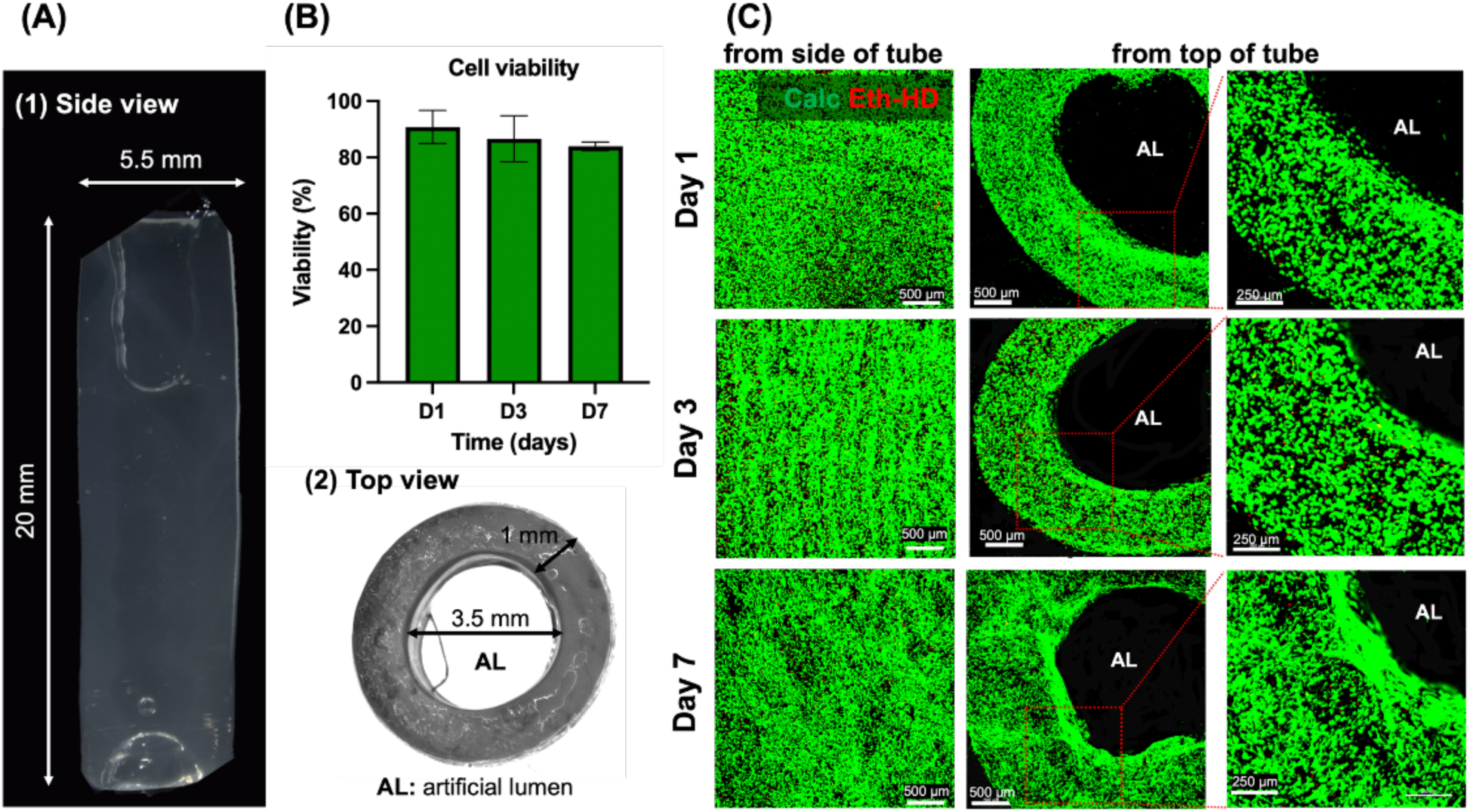
Assessment of printed tubular constructs and cell viability. **(A1)** Side view and **(A2)** top view of representative printed tubes (length 1.5 cm, inner diameter 4.6 mm). **(B)** Quantification of cell viability within the FGXC hydrogel formulation at culture days 1, 3, and 7. Cell viability data were obtained from independently stained 9 different printed tubes. Error bars represent standard deviations. Statistical analysis (Student T test) was performed using Prism5 Software. **(C)** Representative cell viability staining by Calcein-AM (green, viable cells) and Ethidium Homodimer (Eth-HD; red, dead cells) after printing at days 1, 3, and 7 of (B).

At day 3, scattered endothelial clusters as well as punctual CD31^+^ staining appeared along the artificial lumen indicating the onset of endothelial differentiation and the partial lining of the luminal surface by endothelial cells **(Fig. 5A1).** By day 7, CD31^+^ cells elongated and aligned circumferentially along the artificial lumen **(Fig. 5A2, 5B2 and 6A2)**. This was accompanied by emerging cord-like vascular structures within the surrounding mesoderm-like tissue structure. By day 10, the cord-like structures evolved into an interconnected CD31^+^ capillary-like network with early branching and sprouting morphologies, both characteristics of vasculogenic and angiogenic processes **(Fig.5A3).** These findings demonstrate that bioprinted hiMPCs rapidly differentiate into endothelial lineages that not only line the luminal surface of the artificial tube but also self-organize into vascular-like networks, thereby recapitulating key steps of developmental vessel formation *in vitro*. Consistent with these observations, H&E and IF analyses of day 7 constructs revealed an early multilayered vessel wall-like formation lining the lumen of the printed tube along its entire circumference **(Fig. 5B1-B2 and 6A2 & A6)**. The lumen was lined by flattened CD31⁺ endothelial cells resembling an intima-like layer, which was encased by a circumferential layer composed of multiple sheets of αSMA⁺ smooth muscle cells (SMCs) forming a nascent tunica media. The vessel wall-like structure was surrounded by a broad, highly vascularized and loosely organized mesodermal compartment containing CD150^+^, CD44^+^, CD45^+^ and CD34^+^ hematopoietic as well as vascular progenitors closely mimicking the adventitial layer of native blood vessels **(Fig. 5B3–B5)** (Campagnolo et al., 2010; Davis & Senger, 2005; Ergun et al., 2008; Kleefeldt et al., 2022; Worsdorfer et al., 2017; Zengin et al., 2006). Collagen IV deposition, predominantly underlying the endothelial layer lining the luminal surface of the printed tube **(Fig. 5B6),** indicated early basement membrane-like formation and extracellular matrix remodeling conducive to the development of a multilayered vessel wall (Bahramsoltani et al., 2014; Gross et al., 2021; Kanie et al., 2012; Mak & Mei, 2017). Collectively, these findings demonstrate that the printed hiMPCs differentiate into all major vascular wall cell types, including endothelial, mural and progenitor lineages within one week of culture. At later time points this early vessel-like structure undergoes progressive maturation and remodeling into a multilayered wall construct surrounding a lumen with a diameter up to 1-1.5 mm.

**Figure 5:**
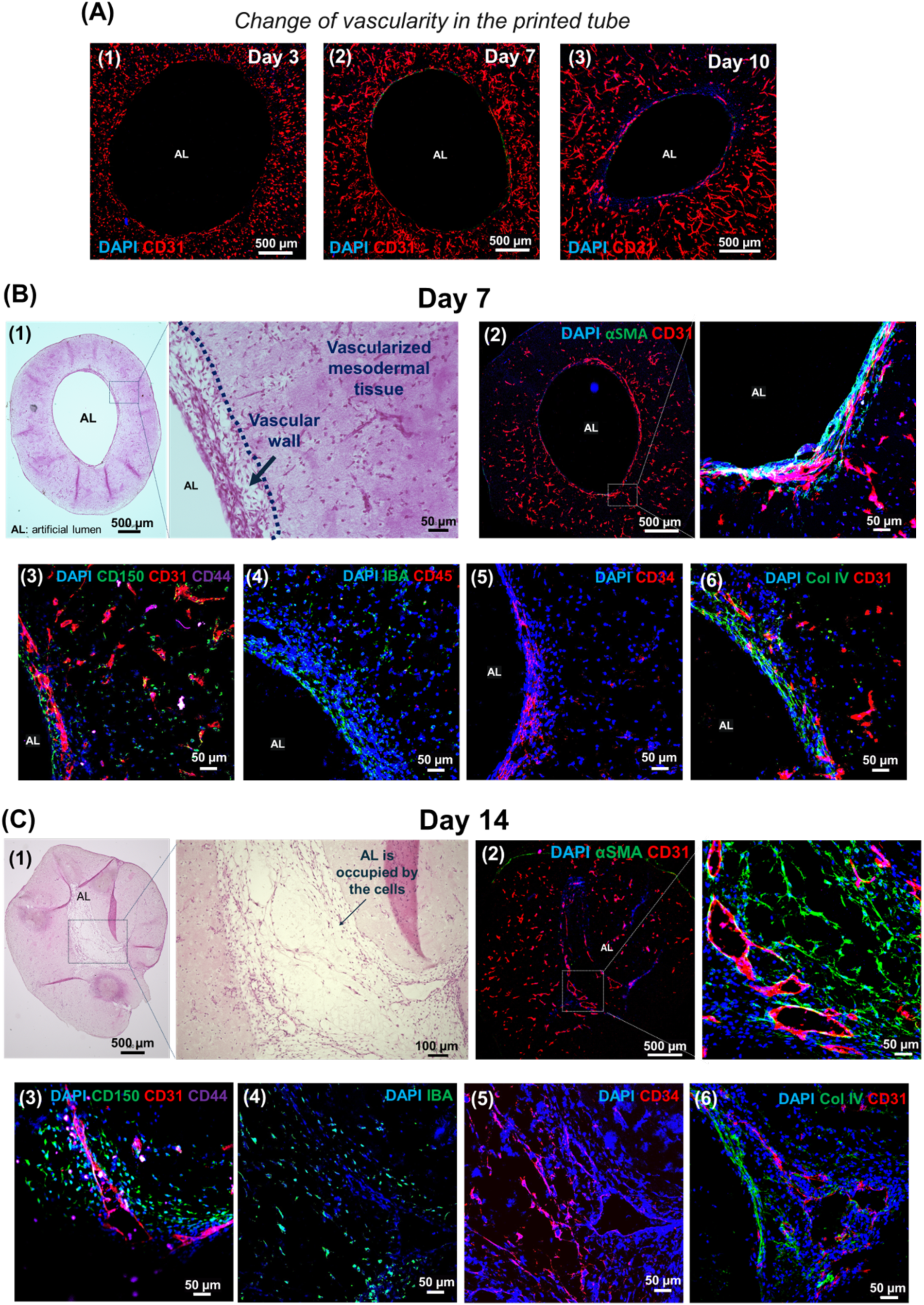
Biological evaluation of hiMPC-laden FGXC printed tubular constructs. **(A)** IF staining demonstrates progression of vascular network formation from d3 to d10. At day 3 **(A1)**, CD31 staining (red) appears sparse and discontinuous, predominantly presenting as dot-like signals, with minimal formation of endothelial cell cords or networks. By day 10 **(A3)**, elongated CD31^+^ endothelial cells lining the artificial lumen become evident, accompanied by the emergence of capillary-like cell cords and an interconnected, branching vascular network. **(B)** Morphology and cellular differentiation at d7. (B1) H&E staining of 10 µm sections reveals an immature vascular morphology characterized by a relatively thin vessel wall surrounded by mesoderm-like tissue. **(B2)** CD31^+^ endothelial cells (red) line the artificial lumen (AL), while αSMA^+^ smooth muscle-like cells (green) form an early media-like layer. **(B3–B5)** Cells expressing hematopoietic progenitor cell markers (CD150, CD44 and CD45) as well as Iba1^+^ macrophage-like cells are present within the wall of the tubular constructs. **(B6)** Collagen IV (green) deposition indicates early extracellular matrix and basement membrane–like formation, particularly within the intima-like region of the tube wall. **(C)** Advanced structural organization at day 14. **(C1)** Vascular constructs show partial narrowing of the AL (H&E). **(C2)** IF images reveal CD31^+^ vascular structures within the former AL that are lined by CD31^+^ endothelial cells (red) and covered by αSMA^+^ pericyte- or smooth muscle–like cells (green). **(C3–C6)** At day 14, cells expressing hematopoietic progenitor cell markers (CD150, CD44, CD34), as well as Iba1^+^ macrophage-like cells are still present within the vascular constructs. Collagen IV is prominently deposited around the CD31^+^ endothelial lining of mid-sized and microvascular structures.

**Figure 6:**
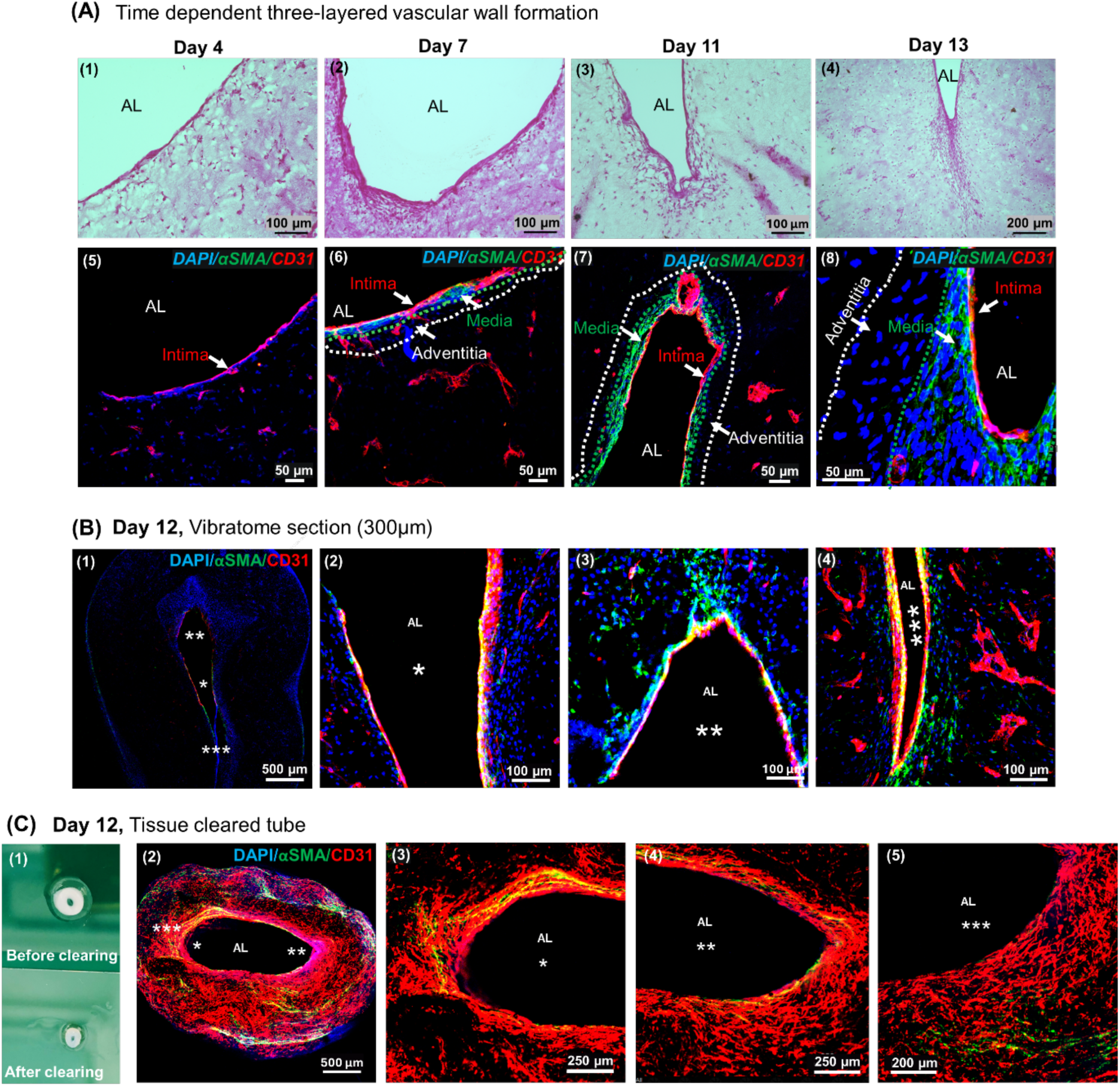
Progressive formation of a multilayered vascular wall over time. H&E **(A1–A4)** and IF **(A5–A8)** stainings demonstrate enhanced vascular wall organization between d4 and d13. CD31^+^ ECs line the artificial lumen (AL) as early as day 4. From day 7 onward, αSMA^+^ peri-endothelial cells (PECs, green) are located in close proximity to the endothelial layer. This indicates the emergence of a nascent media-like compartment that progressively develops into two or more concentric layers **(A6–A8)**. Dotted white lines delineate the transition to the outer adventitia-like layer. **(B)** IF staining for CD31 and αSMA in 300 µm vibratome sections illustrates the spatial arrangement of endothelial and peri-endothelial layers within the tubular constructs. **(C1)** Representative images of printed tube (day 12) before and after tissue clearing. **(C2-C5)** Maximum intensity projection of a z-stack at lower **(C2)** and at higher magnifications of regions indicated by asterisks in C2 **(C3-C5)** highlighting the spatial organization of endothelial and peri-endothelial cells within the constructs.

By day 14, H&E and immunofluorescence staining of tissue sections obtained from such constructs showed partial narrowing of the artificial lumen **(Fig. 5C1 & 6A4)** while they retained their well-organized, multilayered vessel wall-like structure consisting of a CD31^+^ endothelial intima, an αSMA^+^ medial layer and a relatively thick adventitial compartment populated by progenitor cells and IBA1^+^ macrophage-like cells **(Fig. 5C2–C6)**. H&E staining of cultured printed constructs at successive time points (d4, d7, d11, and d13) demonstrated an initial formation of a cellular layer surrounding the artificial lumen **(Fig. 6A1)**. This was followed by progressive cell deposition around the emerging intima-like layer, leading to the development of a media-like compartment and further structural maturation as indicated by the formation of a multilayered vessel wall and an interconnected vascular network (**Fig. 6A2–A4**). These observations were confirmed by IF shown in **Fig. 6A5–A8**. To visualize the three-dimensional organization of these developing vascular networks, day-12 FGXC constructs were subjected to tissue clearing and whole-mount IF for CD31 and αSMA **(Fig. 6B-C)**. The lumen was lined by a continuous CD31^+^ endothelial monolayer, surrounded by αSMA^+^ mural cells and an outer loosely organized adventitia-like layer that additionally harbored progenitor cells. Together this resembled the intima-media-adventitia architecture of the native vascular wall. Maximum-intensity projections further revealed densely branched microvessels extending from the inner wall of the central “mother vessel” into the surrounding mesoderm-like tissue compartment containing undifferentiated or less-differentiated hiMPCs. These observations demonstrate that hiMPC-derived cells can self-organize into an interconnected complex vasculature spanning multiple scales, ranging from potentially perfusable macrovessels with multilayered wall architectures to capillary-like microvessels within the bioprinted tubes with a luminal diameter exceeding 1 mm **(Fig. 6C).**

### Integration of Prevascularized Mesodermal Organoids with Bioprinted “Mother Vessels”

To extend vascularization beyond the printed tube wall, prevascularized mesodermal organoids (Schmidt et al., 2022) were generated and co-cultured with the bioprinted FGXC “mother vessels”. The organoids were generated from hiPSCs in custom made agarose microwell molds. Due to this standardized production process, organoids displayed a highly similar size and vascularization pattern **(Fig. 7A-B).** Histological analysis revealed that the organoids consisted of a loosely arranged mesenchymal core surrounded by a more compact peripheral cell layer **(Fig. 7B2)**. Whole-mount immunofluorescence analyses confirmed an extensive CD31^+^ endothelial network within the organoids **(Fig. 7B3**).

**Figure 7:**
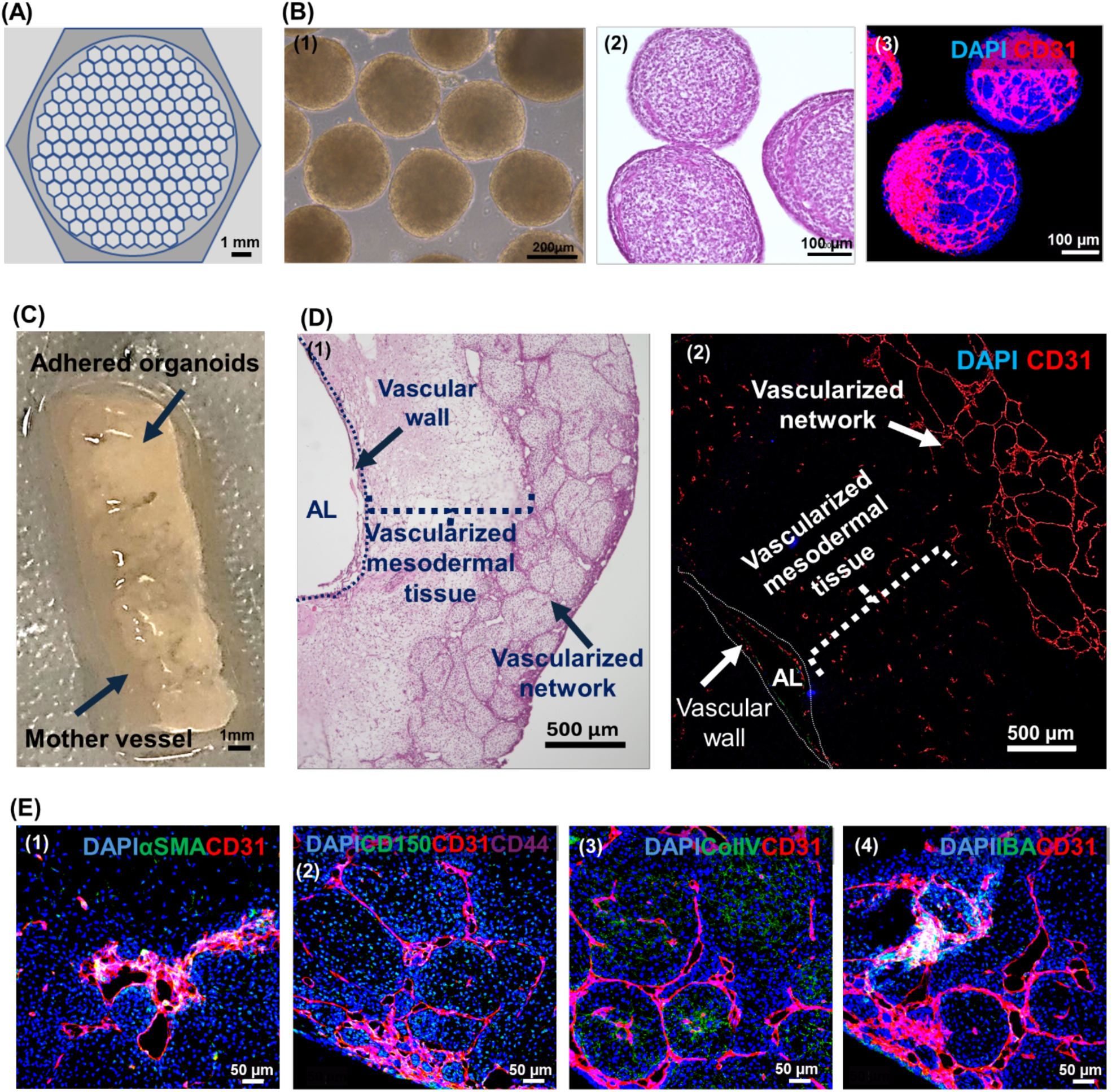
Integration of vascularized mesodermal organoids with biofabricated vascular constructs. **(A)** Schematic illustration of the agarose micro-molds used for mesodermal organoid formation. **(B)** Characterization of mesodermal organoids at d7. Bright-field **(B1)** and H&E images **(B2)** reveal mesodermal organoids with uniform size and morphology that lack central cavitation. The mesodermal organoids are characterized by a dense CD31^+^ vascular network **(B3)**. **(C1)** Prevascularized mesodermal organoids of **B** are incorporated onto the biofabricated vascular tube (length: 2 cm, day 4 construct). **(D)** After three days of co-culture, the organoids fused with each other and cover the outer surface of the vascular construct. **(E1-4)** The covering organoid layer forms an interconnected vascular network with surrounding αSMA^+^ and Iba1^+^ cells as well as Collagen IV deposition indicating early extracellular matrix and basement membrane-like formation.

Approximately 750 organoids were subsequently co-cultured with each 4 day-old bioprinted “mother vessel” for 3 days **(Fig. 7C).** The organoids not just simply adhered to the outer surface of the printed tubes but gradually fused with each other as well as with the “mother vessel” wall. This formed a continuous tissue-like covering of the “mother vessel” **(Fig. 7D).**

Vascular networks from adjacent organoids fused and moreover showed spatial continuity with the network inside the mesoderm-like tissue compartment surrounding the bioprinted “mother vessel,” indicating the onset of structural but not yet functional connection **(Fig. 7E1).** Histological and IF analyses showed vascular organization and maturation within the fused constructs. CD31^+^ endothelial cells and αSMA^+^ mural cells were distributed throughout the organoid-derived layer, forming vessel-like structures comparable in organization to those within the printed “mother vessel” **(Fig. 7E1–E2)**. Collagen IV deposition along vessel walls **(Fig. 7E3)** indicated early basement membrane-like formation as a marker for maturation. Furthermore, appearance of Iba1^+^ macrophage-like cells **(Fig. 7E4)** suggested immune cell presence that is crucial for vascular remodeling *in vivo*.

As a proof-of-concept, the perfusability of the mother vessel was evaluated in a bioreactor using a pulsatile flow generated by a pump connected to the perfusion chamber. The mother vessel was clamped at both ends to the inlet and outlet tubing of the reactor (Fig. S3A, Video). Basal medium was perfused through the system, with pulsatile flow evident at higher magnification in the video recording (Fig. S3B, Video). Importantly, higher magnification analysis revealed no detectable leakiness along the length of the mother vessel. Only a minor amount of the fluid entered the chamber at the clamped inflow end (Fig. S3C, Video), indicating high stability and tightness of the mother vessel wall and supporting its suitability for perfusion.

## Conclusion

In this study, we successfully established the bioprinting of centimeter-scale “mother vessel” constructs using hiMPCs and a newly optimized bioink composed of piscine GelMA, procine GelMA, xanthan gum, and fibrillar Col I. This formulation provided both printability and biological compatibility enabling the single-step generation of centimeter-scale macrovessels that we termed “mother vessels”. In contrast to most previous approaches that were restricted to microvascular networks or simple endothelialized channels (Wu et al., 2024), the printed vessel constructs described here closely resembled macrovessels, approaching the realistic dimensions of native human vessels in the centimeter range, and spontaneously developed key structural hallmarks of native vessel walls within just a few days after printing.

Within just one week, hiMPCs underwent differentiation into vascular cell types and self-organized into early endothelial and mural layers reminiscent of embryonic vascular development. The constructs exhibited a characteristic tri-layered organization consisting of a luminal CD31^+^ endothelial lining, an intermediate αSMA⁺ medial layer and a loosely organized adventitia-like outer layer which also harbored CD150^+^/CD45^+^/CD34^+^ progenitor cells. This vessel wall-like structure was surrounded by a thick vascularized compartment also containing CD150^+^/CD45^+^/CD34^+^ progenitor cells that are considered to differentiate into both hematopoietic and endothelial cells. This pattern is similar to early embryonic vessel formation (Siekmann, 2023). Strikingly, IBA1^+^ macrophage-like cells appeared despite their absence from the initial cell suspension. This finding indicates that the vascular constructs enable intrinsic differentiation of hiMPCs not only into vascular lineages but also non-vascular cell types, an aspect that will be investigated in detail in future studies. This is of particular importance since macrophages orchestrate angiogenesis and vascular remodeling, e.g. by release of proangiogenic mediators such as VEGF. Furthermore, macrophages are critical for tissue homeostasis as well as plasticity. This intrinsic driven spontaneous differentiation of different cell types from a single progenitor source distinguishes this system from most engineered vascular models today, which mainly rely on differentiation into different cell types prior bioprinting (Dell et al., 2025; Kong & Wang, 2023).

To mimic prevascularized tissue modules and advance the concept of modular vascularization, prevascularized mesodermal organoids were co-cultured with the bioprinted “mother vessels”. The organoids contained intrinsic microvascular networks and gradually fused with the outer “mother vessel” wall forming a continuous, vascularized tissue envelope. Microvessels extending from the organoids toward the bioprinted wall indicated early structural but not yet functional interconnection. This demonstrates that prevascularized organoids may serve as biological building blocks. In combination with bioprinted large “mother vessels” as conduits, this may promote formation of hierarchically organized vascular networks capable of supplying larger engineered tissue constructs. Controlled perfusion may not only promote further functional vascular integration between the “mother vessel” and the surrounding fused organoids but also counteract lumen narrowing of the printed “mother vessels”. Furthermore, these biomechanical stimuli are expected to enhance vascular maturation, resembling the effects induced by the onset of the heartbeat during embryonic development (Laowpanitchakorn et al., 2024).

Taken together, this study presents a bioprinting strategy that enables the generation of centimeter-scale, self-organizing vascular tubes directly from hiMPCs in a single bioprinting step. The optimized bioink supports high viability, spatial organization and intrinsic cell differentiation into vascular but also non-vascular cells reflecting key processes of embryonic vessel development. Integrating these printed “mother vessels” with prevascularized organoids establishes a modular system for hierarchical vascularization of large tissue constructs. In this system, macrovascular conduits enable vascular integration with the host, while microvascular networks self-assemble within the tissue to provide nutrient and oxygen supply, thereby addressing one of the major obstacles in tissue biofabrication. Using the newly designed perfusion chamber system, the mother vessel was successfully perfused under pulsatile basal medium flow without detectable leakage while clamped within the system. Together with pre-vascularized organoid-mother vessel construct, the demonstrated perfusability of the mother vessel supports its suitability for establishing an *in vitro* circulatory system capable of supplying complex tissue constructs at large scale (Figure 8). Moreover, this platform would enable tissue- and organ specific analyses at molecular, cellular, and humoral levels.

**Figure 8:**
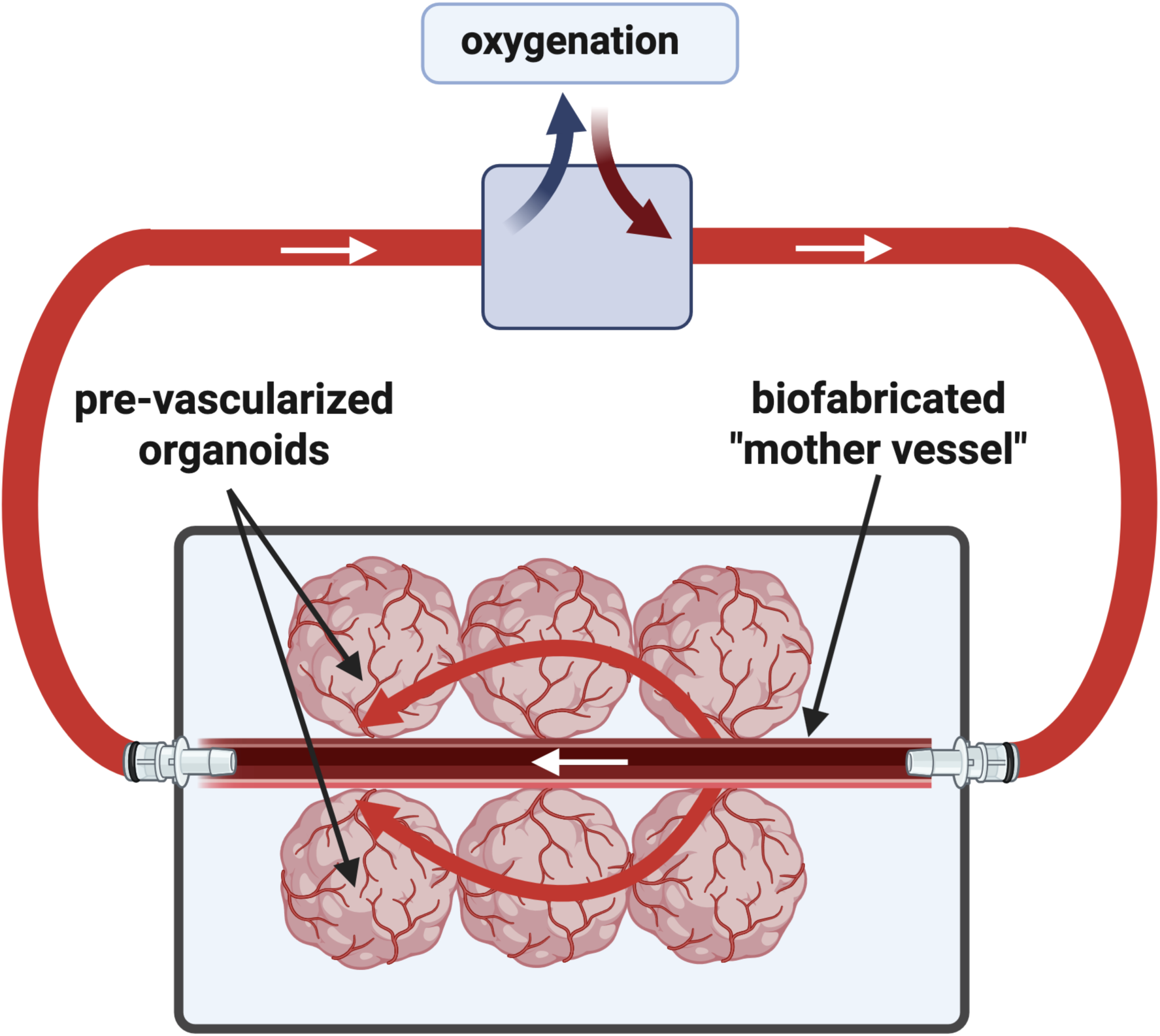
This graphical representation illustrates how the pre-vascularized organoids are connected to the mother vessel via their own intrinsic vascular network, as demonstrated in this manuscript. In addition, it provides a future perspective on how such tissue constructs could be integrated into a perfusion system (as shown using a perfusion chamber in the proof-of-concept video in Supplementary Video S3). This approach enables the establishment of large-scale vascularized tissue models that are suitable for experimental perfusion through mother vessel at the macrovascular level.

## Acknowledgments

This research was funded by the Deutsche Forschungsgemeinschaft (DFG, German Research Foundation; Project No. 326998133) within the framework of the Collaborative Research Centre Transregio 225 (TRR 225; subproject B04; funding period 2022-2025: Prof. Süleyman Ergün and Prof. Jürgen Groll; funding period 2026-2029: Prof. Florian Kleefeldt and Prof. Jürgen Groll). We thank Dr. Taufiq Ahmad for providing the 3D-printed microwell molds, Dr. Philipp Stahlhut for cryo-SEM imaging, and Csaba Gergely for porcine GelMA and piscine GelMA synthesis (all: Department for Functional Materials in Medicine and Dentistry, University of Würzburg).

## Supplementary Information

**Figure S1:**
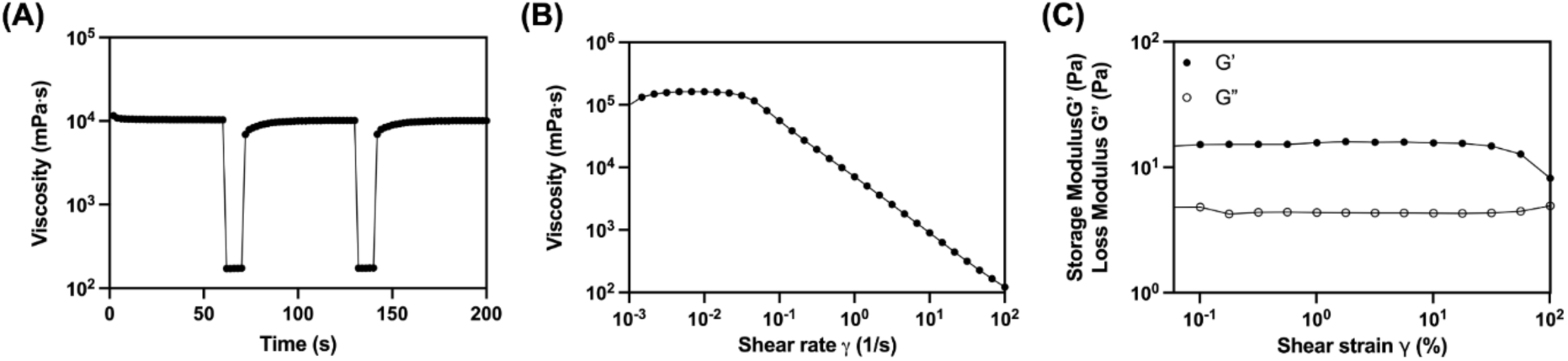
Rheological characterization of the XG-based embedding medium. **(A)** Shear stress recovery over time. **(B)** Flow curve of viscosity as a function of shear rate. **(C)** Frequency sweep test showing the gel-like and viscoelastic liquid-like behavior of the support bath.

**Figure S2:**
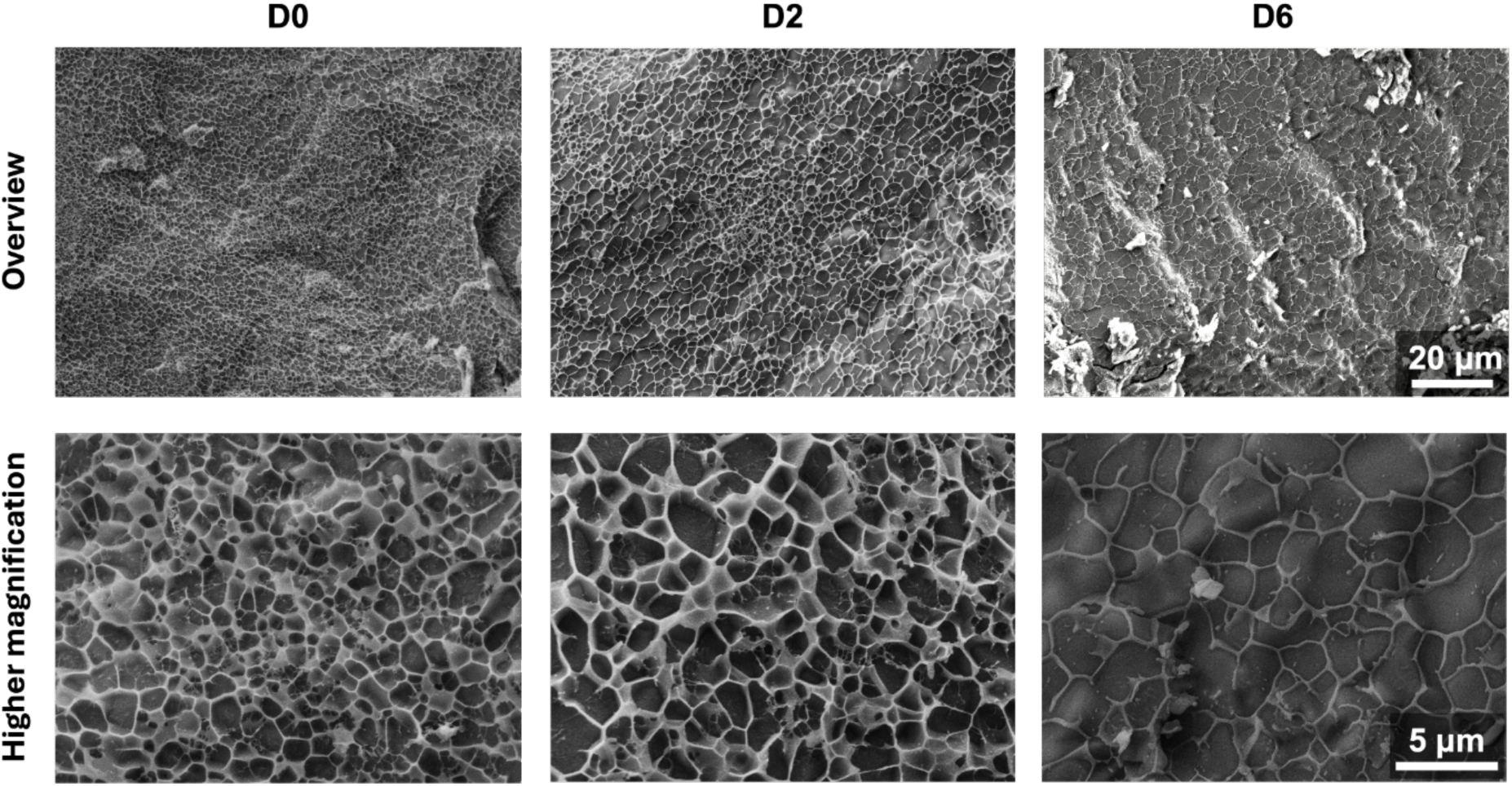
Microstructural changes in FGXC overtime. FGXC samples without cells were incubated in PBS for 6 days. Changes in the polymeric network density were observed, suggesting swelling of the hydrogel and release of XG over time.

**Figure S3A:**
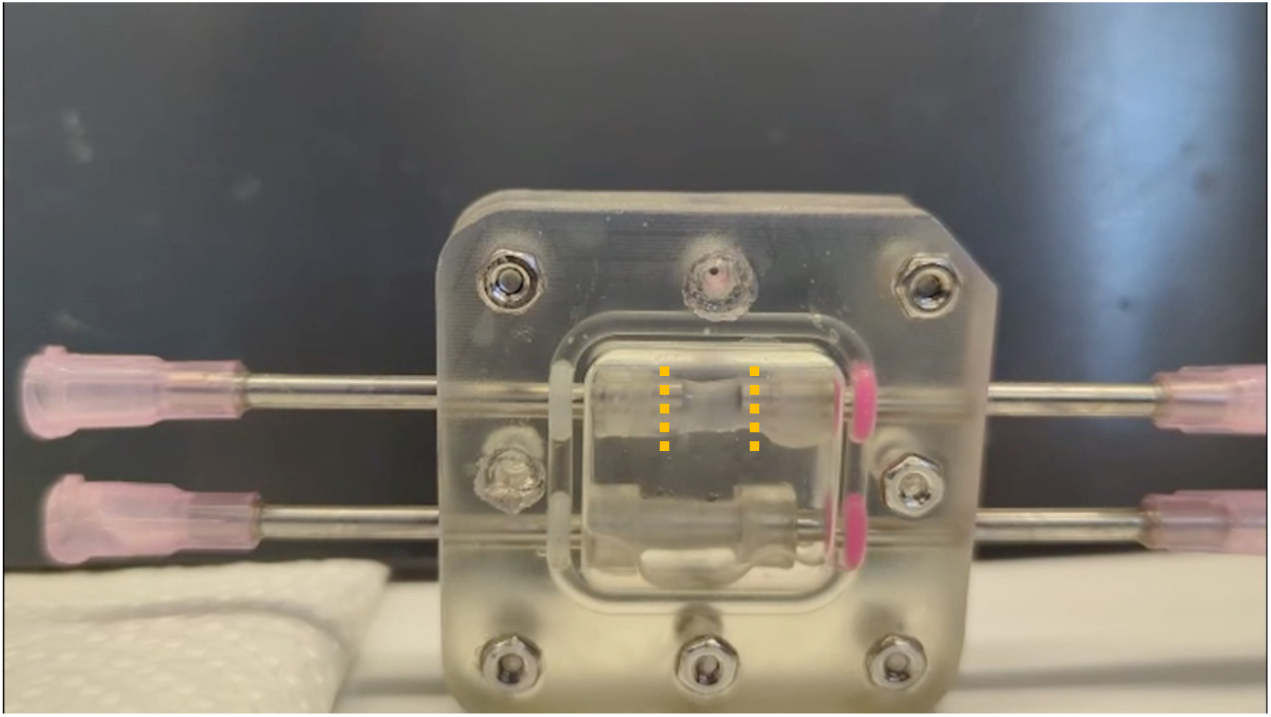
This video demonstrates perfusion of the mother vessel tube, which is clamped at both ends (indicated by doted orange lines) to the inlet and outlet openings of the metal tubing. Perfusion was achieved using a peristaltic pump, with basal culture medium flowing through the construct containing the mother vessel and exiting at the open end of the tubing as visible in the video.

**Figure S3B:**
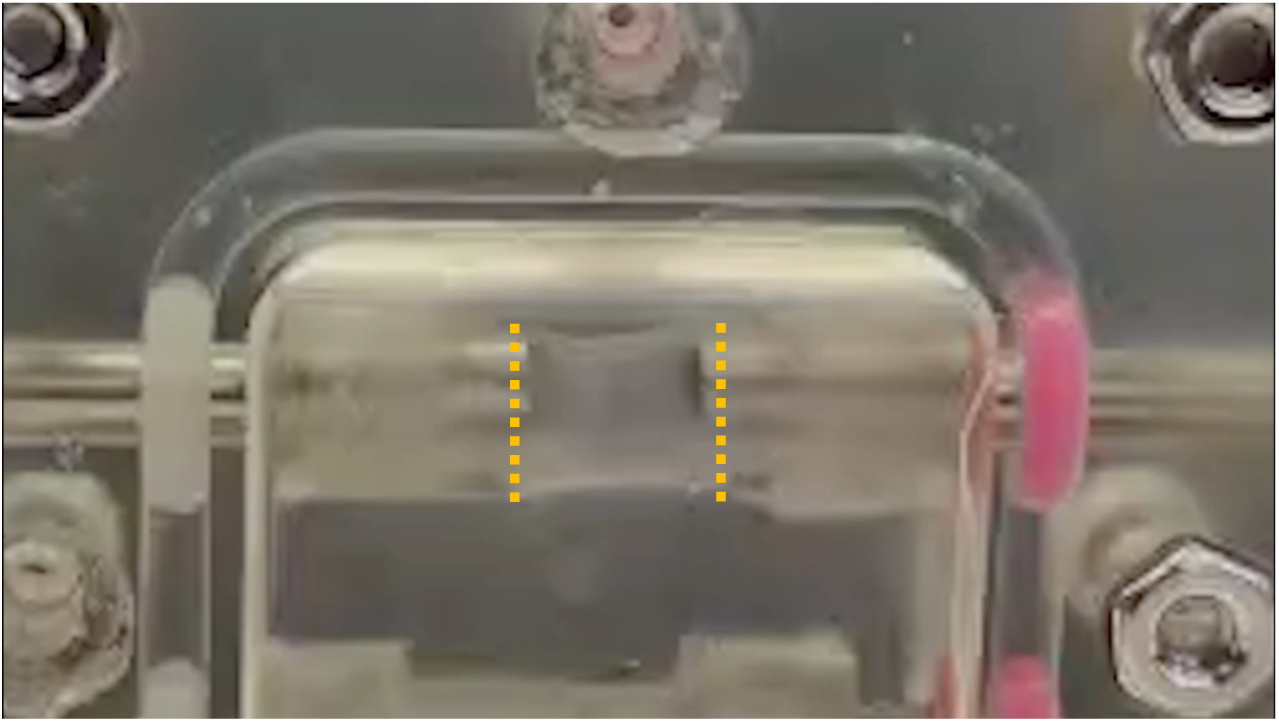
This video recording in higher magnification demonstrates the pulsatile perfusion of the mother vessel tube within the section marked by doted lines.

**Figure S3C:**
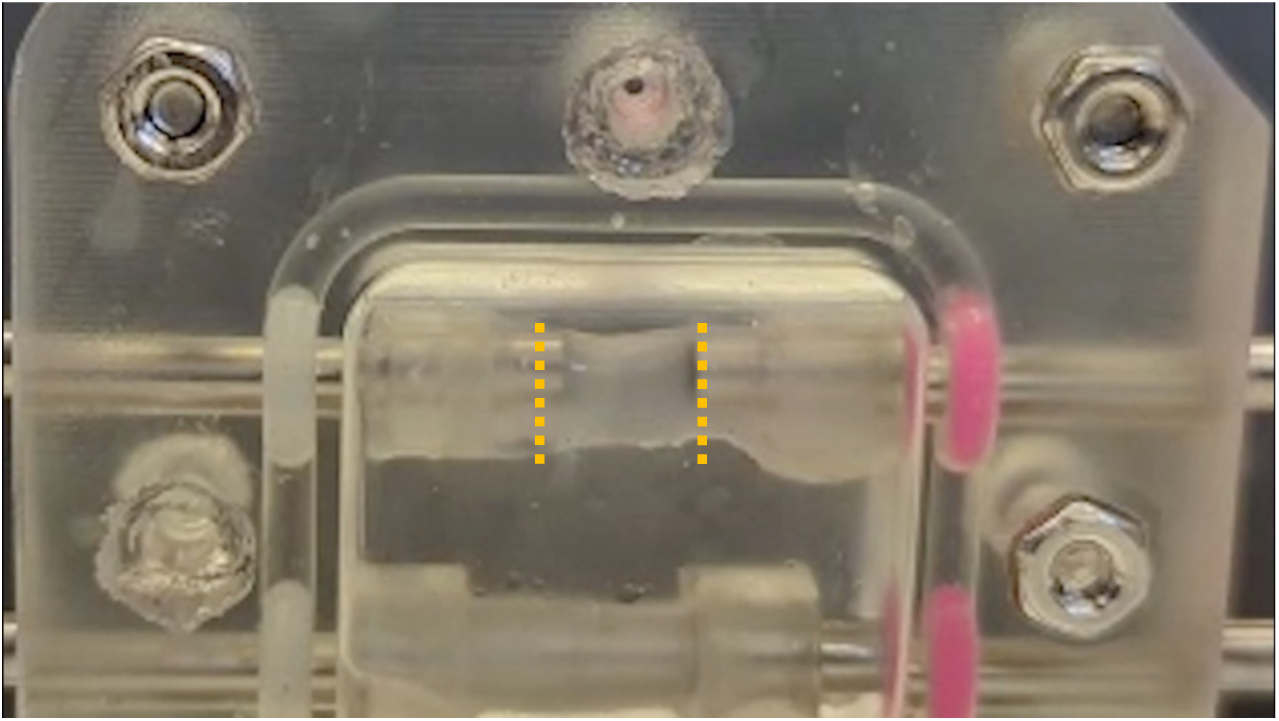
This high-magnification video recording demonstrates the absence of detectable leakage through the mother vessel wall.

