## Supplementary figures and images for "Generation of Self-Organising Macrovascular Constructs by Bioprinting human iPSC-Derived Mesodermal Progenitor Cells"

### Figure S1

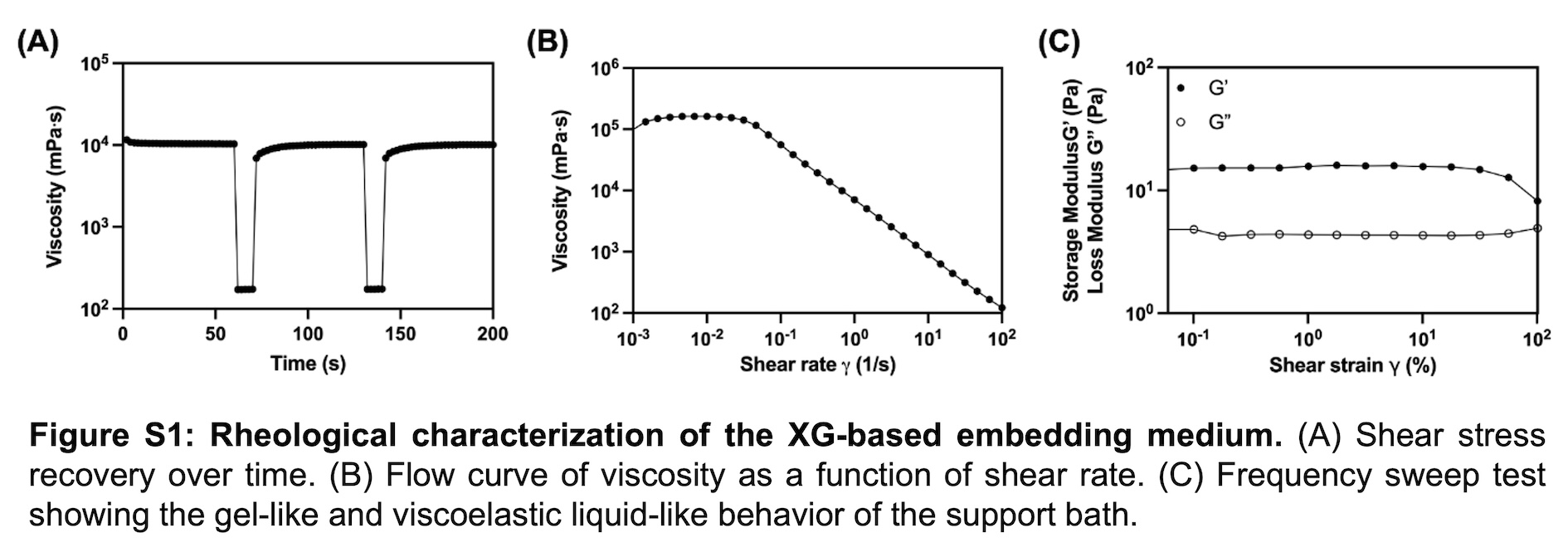

### Figure S2

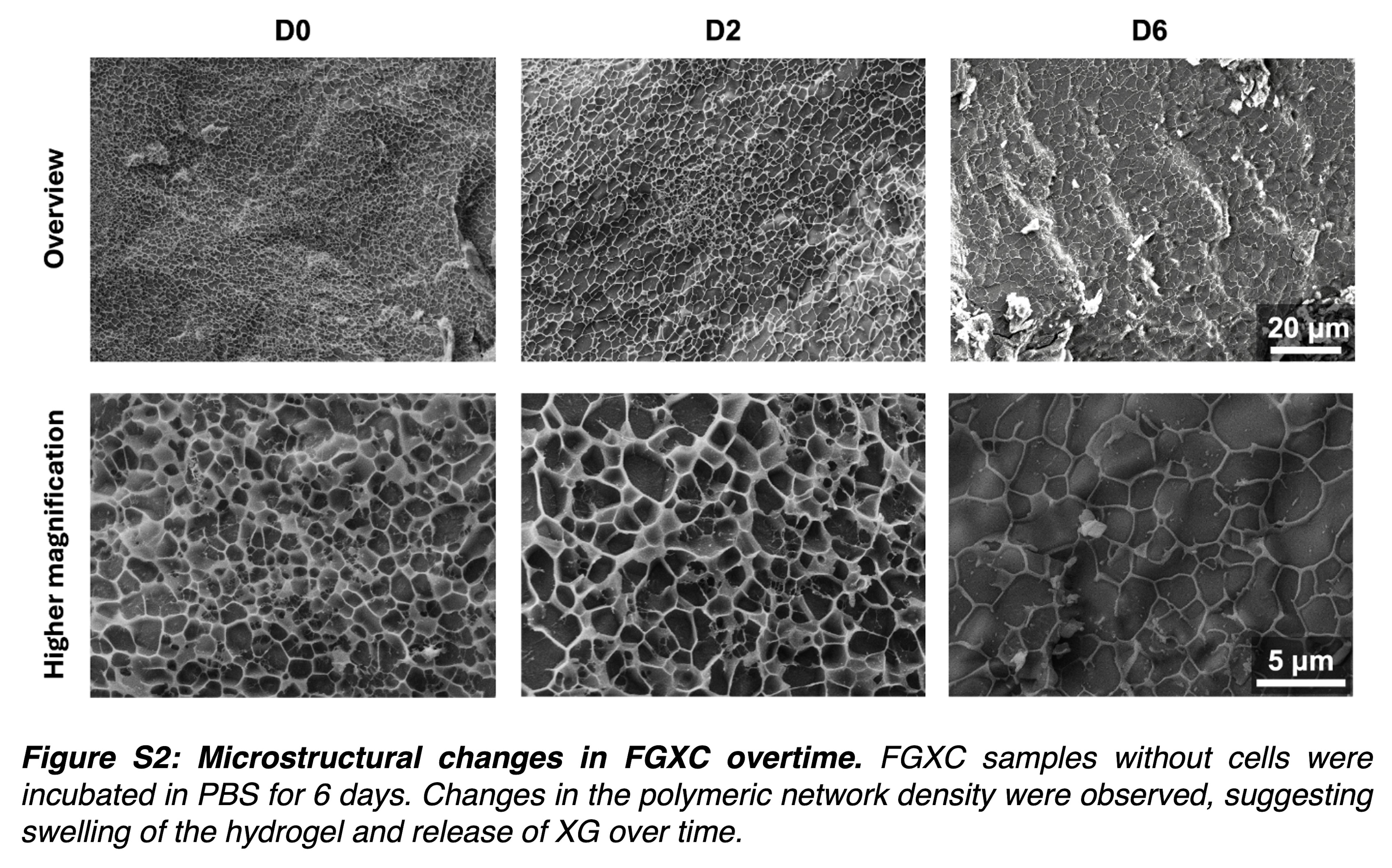

### Figure S3 A

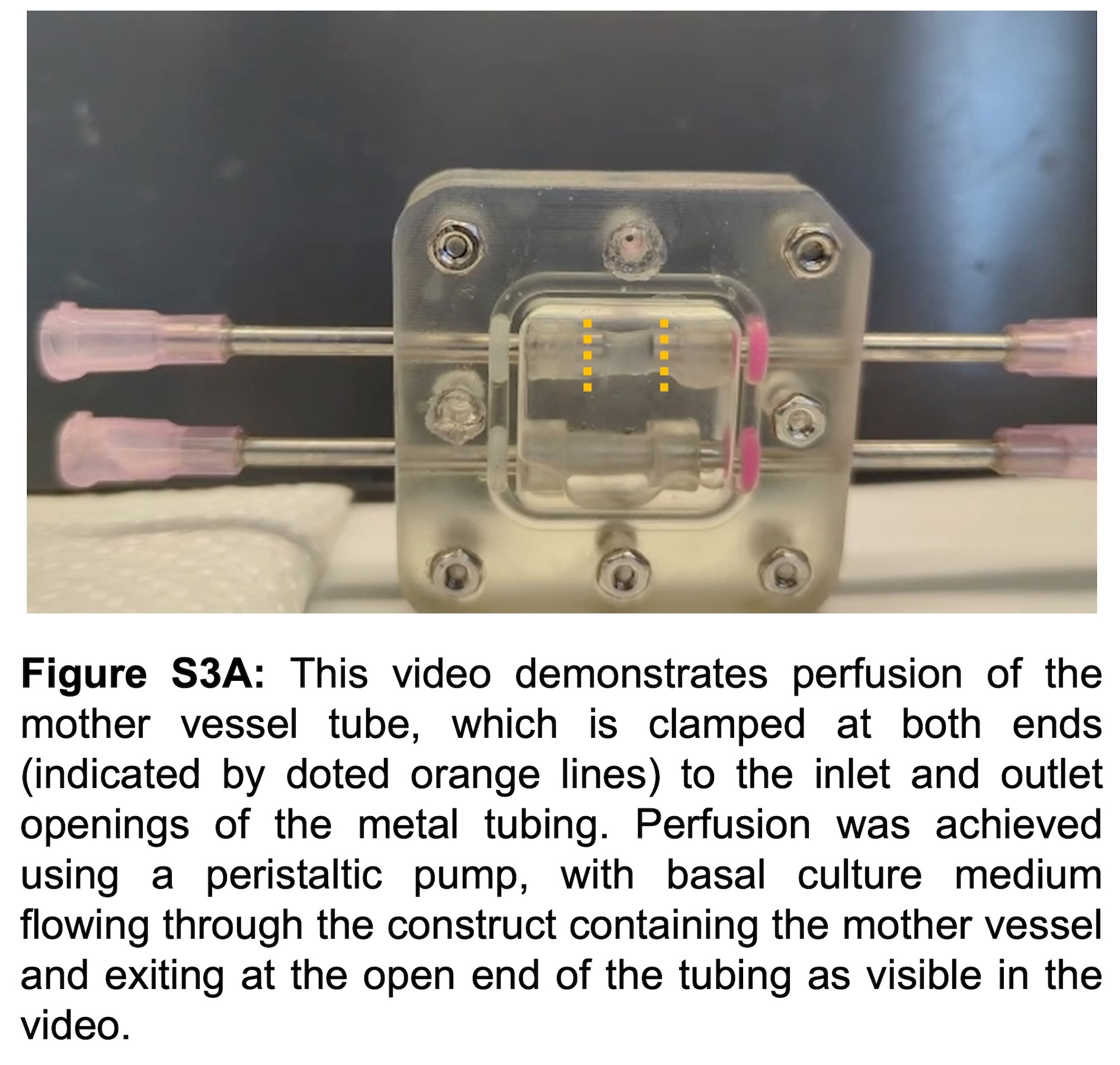

### Figure S3 B

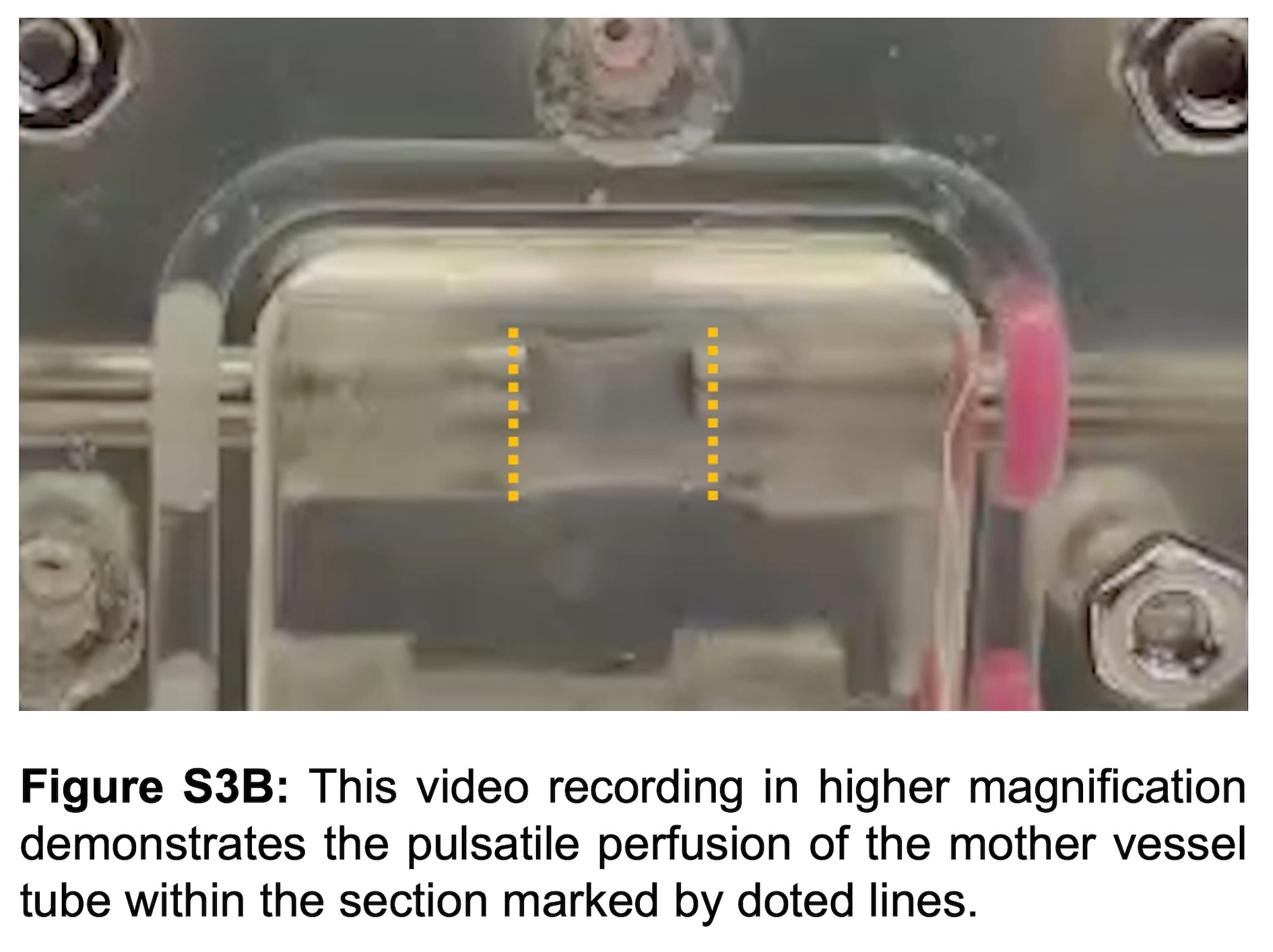

### Figure S3 C

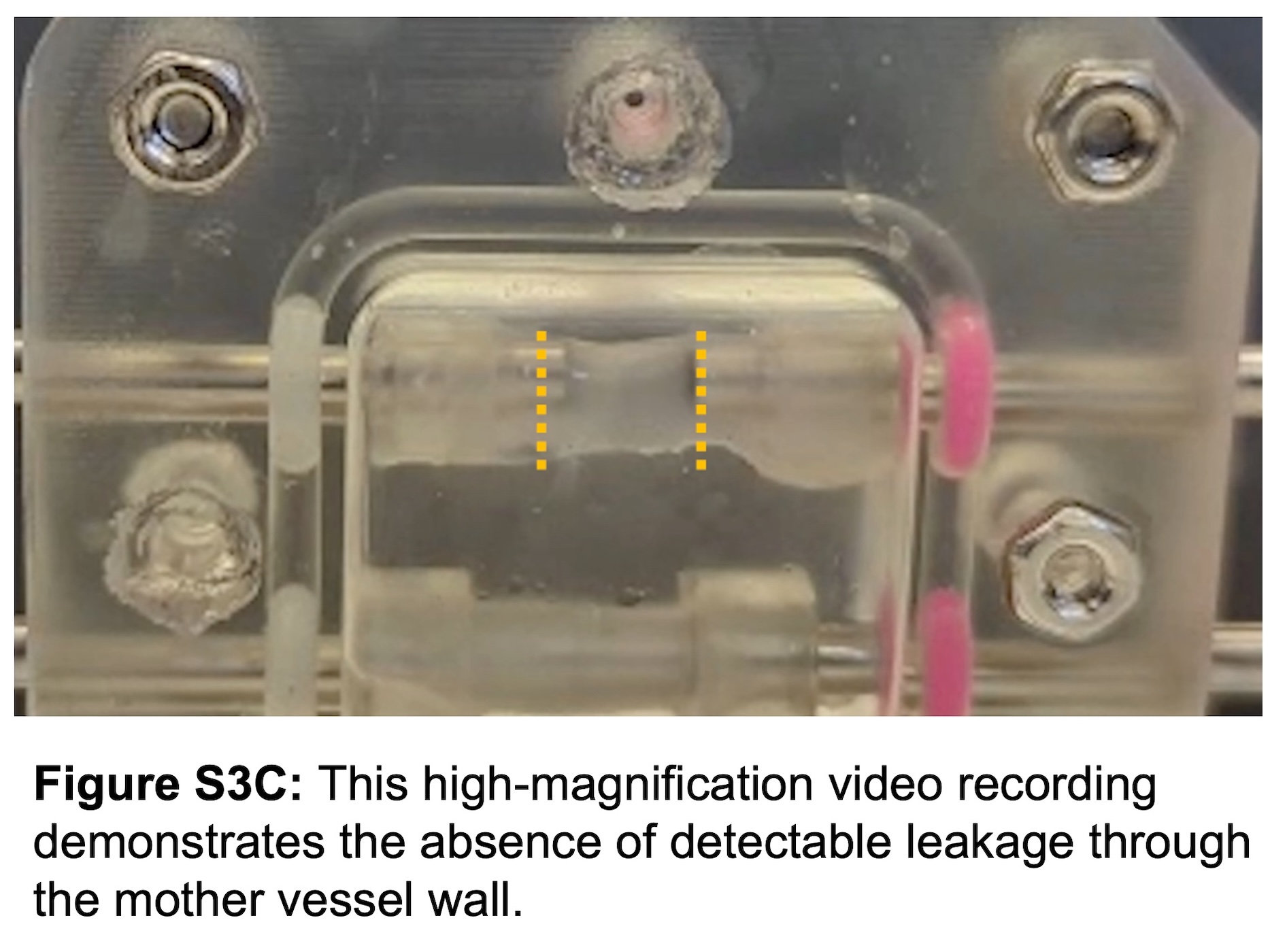
